# Automated measurement of long-term bower behaviors in Lake Malawi cichlids using depth sensing and action recognition

**DOI:** 10.1101/2020.02.27.968511

**Authors:** Zachary V Johnson, Lijiang Long, Junyu Li, Manu Tej Sharma Arrojwala, Vineeth Aljapur, Tyrone Lee, Mark C Lowder, Karen Gu, Tucker J Lancaster, Joseph I Stockert, Jean M Moorman, Rachel L Lecesne, Jeffrey T Streelman, Patrick T McGrath

## Abstract

Measuring naturalistic behaviors in laboratory settings is difficult, and this hinders progress in understanding decision-making in response to ecologically-relevant stimuli. In the wild, many animals manipulate their environment to create architectural constructions, which represent a type of extended phenotype affecting survival and/or reproduction, and these behaviors are excellent models of goal-directed decision-making. Here, we describe an automated system for measuring bower construction in Lake Malawi cichlid fishes, whereby males construct sand structures to attract mates through the accumulated actions of thousands of individual sand manipulation decisions over the course of many days. The system integrates two orthogonal methods, depth sensing and action recognition, to simultaneously measure the developing bower structure and classify the sand manipulation decisions through which it is constructed. We show that action recognition accurately (>85%) classifies ten sand manipulation behaviors across three different species and distinguishes between scooping and spitting events that occur during bower construction versus feeding. Registration of depth and video data streams enables topographical mapping of these behaviors onto a dynamic 3D sand surface. The hardware required for this setup is inexpensive (<$250 per setup), allowing for the simultaneous recording from many independent aquariums. We further show that bower construction behaviors are non-uniform in time, non-uniform in space, and spatially repeatable across trials. We also quantify a unique behavioral phenotype in interspecies hybrids, wherein males sequentially express both phenotypes of behaviorally-divergent parental species. Our work demonstrates that simultaneously tracking both structure and behavior provides an integrated picture of long-term goal-directed decision-making in a naturalistic, dynamic, and social environment.

## 1. INTRODUCTION

Natural behaviors are often expressed over long timescales. For example, many construction, navigation, hunting/foraging, and social behaviors are executed over timescales ranging from many hours to weeks and are critical for survival and reproduction in a wide range of invertebrate and vertebrate species (Tucker 1981, Feng, Fergus et al. 2015, Russell, Morrison et al. 2017, Mouritsen 2018). These behaviors may be expressed inflexibly according to fixed sets of rules, or plastically in response to changing environmental and social stimuli. Understanding the underlying logic of long-term behaviors and how they are encoded in the genome and the nervous system will require accurately measuring them as they unfold over extended periods of time in complex, naturalistic, dynamic, and often social environments.

Long-term natural behaviors are also often goal-directed, in which animals integrate external stimuli, internal physiology, and previous experience to coordinate decisions and actions towards a specific goal. For example, many species exhibit construction behaviors in which they manipulate the environment to build extended phenotype structures such as burrows, dens, tunnels, webs, nests, or bowers; and these structures are integral to survival and reproduction (Dawkins 1982, Vollrath 1992, Collias and Collias 2014, Mouritsen 2018). Construction behaviors are particularly excellent models of long-term goal-directed behaviors because the physical structure itself provides a history of an animal’s goal-directed decision-making and also represents a measurable and dynamic external stimulus that continuously modulates decision-making over long timescales. Thus, measuring both the developing structure and the underlying behavioral decisions throughout construction can provide quantitative descriptions of long-term goal-directed decision-making in dynamic environments.

Measuring construction behaviors and other complex natural behaviors in the lab is challenging. Most existing tools for behavioral phenotyping are designed for paradigms in which single test subjects are behaving in simple, static, and often unfamiliar environments with uniform backgrounds over short timescales. In contrast, natural behaviors are often most faithfully expressed over long timescales, in naturalistic environments, and through direct interaction with the environment itself and/or with other individuals. Additionally, during construction behaviors, the individual and/or structure is frequently partially or wholly occluded from view (e.g. subterranean burrows or tunnels, or enclosed nests), making it difficult to measure the developing structure and the underlying behavior. Because of these challenges, natural behaviors are typically quantified through manual observation and scoring, which is labor intensive and limits the potential scope and scale of experimental designs and research questions that can be pursued. Thus, circumventing the need for manual scoring through automated approaches will facilitate investigations of the biological mechanisms regulating natural behaviors.

In this paper we use automated approaches to measure long-term bower construction behaviors in in Lake Malawi cichlids. Lake Malawi is the most species-rich freshwater lake on Earth, home to an estimated 700-1,000 cichlid species that have rapidly evolved in the past 1-2 million years (Kocher 2004). These species vary strongly in many complex traits, including behavior (Kocher 2004, Hulsey, Mims et al. 2010, Maan and Sefc 2013, Johnson, Moore et al. 2019). The high degree of genetic similarity among species (average sequence divergence between species pairs is 0.1-0.25%) (Loh, Bezault et al. 2013, Malinsky, Svardal et al. 2018) enables behaviorally divergent species to be intercrossed in the laboratory to produce hybrids, making Lake Malawi cichlids a powerful system for studying the genetic and neural basis of natural behavioral variation.

About 200 Lake Malawi species exhibit long-term social bower construction behaviors, in which males manipulate sand to construct large courtship structures, or bowers, during mating contexts (York, Patil et al. 2015). Bower behaviors appear to be an example of convergent mating system evolution, mirroring that of Ptilonorhynchidae birds, in which males congregate into leks and construct elaborate bowers for courtship and mating, but not for raising offspring (McKaye, Stauffer et al. 2001). Among bower constructing species in Lake Malawi, two major behavioral phenotypes have repeatedly evolved: “pit-digging,” or construction of crater-like depressions, and “castle-building,” or construction of volcano-like elevations (York, Patil et al. 2015) (**Figure 1**). Both pits and castles are constructed over the course of many days by collecting mouthfuls of sand and spitting the sand into new locations, ultimately giving rise to the final bower structures.

**Figure 1.**
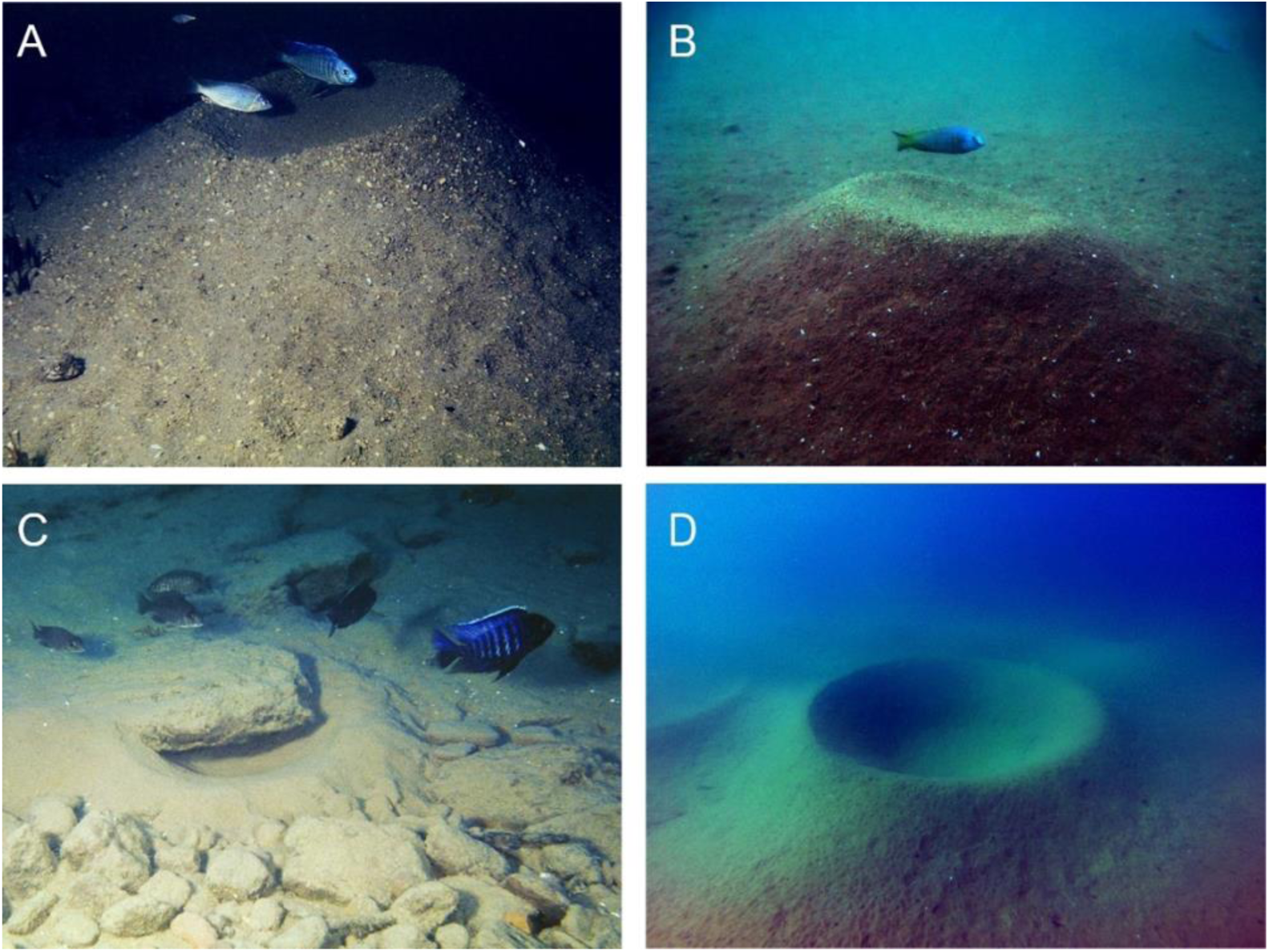
The evolution of bower behaviors in Lake Malawi cichlids. Approximately 200 species of Lake Malawi cichlids exhibit bower behaviors. In these species, sociosexual cues trigger reproductive adult males to construct large courtship structures by manipulating sand with their mouths. The geometric structure of the bower is species-specific. Castles (A,B), or mountain-like elevations, and pits (C,D), or crater-like depressions, are two bower forms that have repeatedly evolved in many species. Some pit-digging species construct pits alongside and partially underneath rocks (C). Photo credit to Dr. Ad Konings (A,C), Dr. Isabel Magalhaes, PhD (B) and Dr. Ryan York (D).

Bower construction behaviors are an excellent opportunity to understand the genetic and neural basis of long-term goal-directed decision-making in a complex and continuously changing environment. However, measuring bower construction in the laboratory is challenging. Bowers are constructed over many days, requiring collection and analysis of large volumes of data. Bowers are constructed in social environments in which multiple individuals can freely interact, making individual tracking difficult. Sand manipulation results in a dynamic background, and the subject male and stimulus females are largely camouflaged against the sand background from a top-down view, both posing difficulties for traditional computer vision strategies. Lastly, scooping and spitting sand during bower construction is behaviorally similar to scooping and spitting sand during feeding, which is performed by both male and female fish over the course of the trial, greatly increasing the difficulty of selectively measuring construction behaviors from video data. In this paper, we integrate two orthogonal methods to automatically track both the bower structure and the thousands of individual sand manipulation decisions made during construction for up to weeks at a time, in multiple species and hybrid crosses, and in many home tank aquariums simultaneously. We use low cost mini computers and video game depth sensors to capture natural species differences in bower form, and we show that a neural network for action recognition accurately classifies bower construction, feeding, and spawning behaviors across hundreds of hours of video data. Through these approaches we gain new insights into bower construction behaviors. We show that distinct behavioral and social contexts emerge over the full course of bower construction, and we show that males (i) construct bowers across many days through punctuated bursts of activity, (ii) construct bowers in spatially repeatable locations across multiple trials, and (iii) exhibit shifts in spatial decision-making during the first days of construction. Additionally, we show that pit-castle F_1_ hybrid males independently express both pit-digging and castle-building behaviors in sequence.

## 2. RESULTS

### 2.1 Assay and recording system for measuring bower behaviors

Lake Malawi bower cichlids construct species-typical bowers in aquariums similar to those observed in the field (York, Patil et al. 2018). However, because bowers are constructed over many days through intermittent bouts of activity, we found that daily 2-3 hour video recordings were insufficient for capturing the behaviors consistently. In order to measure bower behaviors for many days and across many aquariums simultaneously, we collected 10 hours of video data and 24 hours of depth sensing data for 10 days. We used small, inexpensive Raspberry Pi 3 (Pi) computers that could easily be mounted above each tank, and each unit was connected to a small touch screen, an external hard drive for data storage, and an ethernet cord for internet access and interfacing with a common Google spreadsheet file (**Figure 2** and **Figure S1, S3**, and **S4**). For video recording, we connected each unit to a Raspberry Pi camera board that supports HD quality compressed video with a high frame rate; and for depth sensing, we connected each unit to a Microsoft Kinect depth sensor, which has previously been shown to measure distances of natural substrates through shallow creeks (Mankoff and Russo 2013). By optimally positioning the camera and Kinect, we were able to record video and depth data across the sand tray (**Figure 2C**, also see **Figure S5**). For each bower trial, a subject male was introduced to a 50-gallon aquarium containing four adult reproductive females and a sand tray positioned directly beneath the Raspberry Pi camera and Kinect depth sensor for top-down video recording and depth sensing (**Figure 2C** and **Figure S1**).

**Figure 2.**
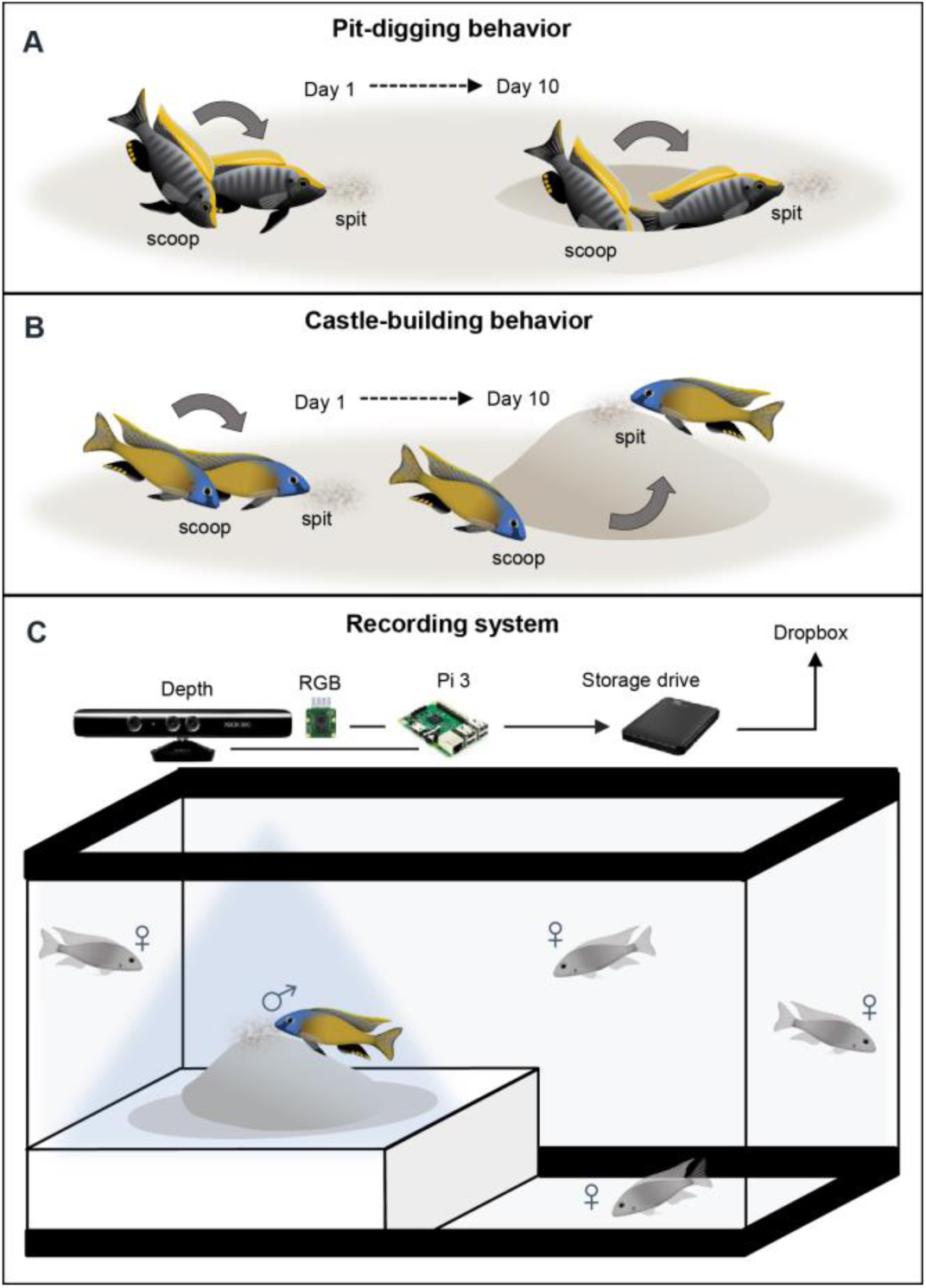
An automated recording system to measure bower behaviors in laboratory aquariums. Bowers are constructed over the course of many days (A,B). Pit-digging involves scooping sand from a concentrated region and spitting it into dispersed locations (A, representation of a *Copidachromis virginalis* male digging a pit). Castle-building involves scooping sand from dispersed locations and spitting it into a concentrated region (B, representation of a *Mchenga conophoros* male building a castle). To measure bower behaviors, we developed a behavioral assay and an automated recording system for standard laboratory aquatics facilities (C). A reproductive adult male is introduced to a 50-gallon aquarium tank containing a sand tray and four reproductive females. The recording system utilizes a Raspberry Pi 3 computer connected to a high-definition RGB camera and a Microsoft Kinect depth sensor for video recording and depth sensing, respectively. Data is stored on an external hard drive and uploaded to Dropbox. The system is remotely controlled by custom Python scripts and a Google documents spreadsheet.

### 3.2 Depth Data

#### 3.2.1 System validation

##### The Kinect detects surface change during bower construction

To validate measurements of depth change, we analyzed the overall volume of sand moved in “bower” trials (in which an experimenter visually identified bowers constructed by the male; n=29 total; pit-digger *Copadichromis virginalis*, CV, n=9; castle-builder *Mchenga conophoros*, MC, n=7; pit-digger *Tramitichromis intermedius*, TI, n=5; pit-castle MCxCV F_1_ hybrid, n=3; pit-castle TIxMCF_1_ hybrid, n=5) and control trials in which no bowers were constructed (n=9 total; CV, n=3; MC, n=3; TI, n=3) by subtracting the initial depth map from the final depth map (for visualization of this calculation see **Figure S5D;** example data shown in **Figure 3A-F**). As a second control, we also analyzed empty tank (no fish) trials to estimate the level of depth change that might be attributed to noise in our depth data. Because males move large volumes of sand during bower construction, we expected to observe larger depth change signals in bower trials compared to control trials. We found that depth change differed strongly between these three conditions, and was much greater in bower building trials (n=29 trials; 1162.7 ± 98.25 cm^3^ volume change) compared to control trials (n=9 trials; 414.5 ± 17.53 cm^3^ volume change) and empty tank trials (n=6 trials; 249.9 ± 24.00 cm^3^ volume change; Kruskal-Wallis χ^2^ = 30.1, p=2.86×10^−7^; **Figure S5**). More information on validation of depth data quality, thresholding, and measurement across timescales is described in the “Validation of Depth Sensing System” section in the Supplementary Materials and Methods, and in **Figures S5A-G**.

**Figure 3.**
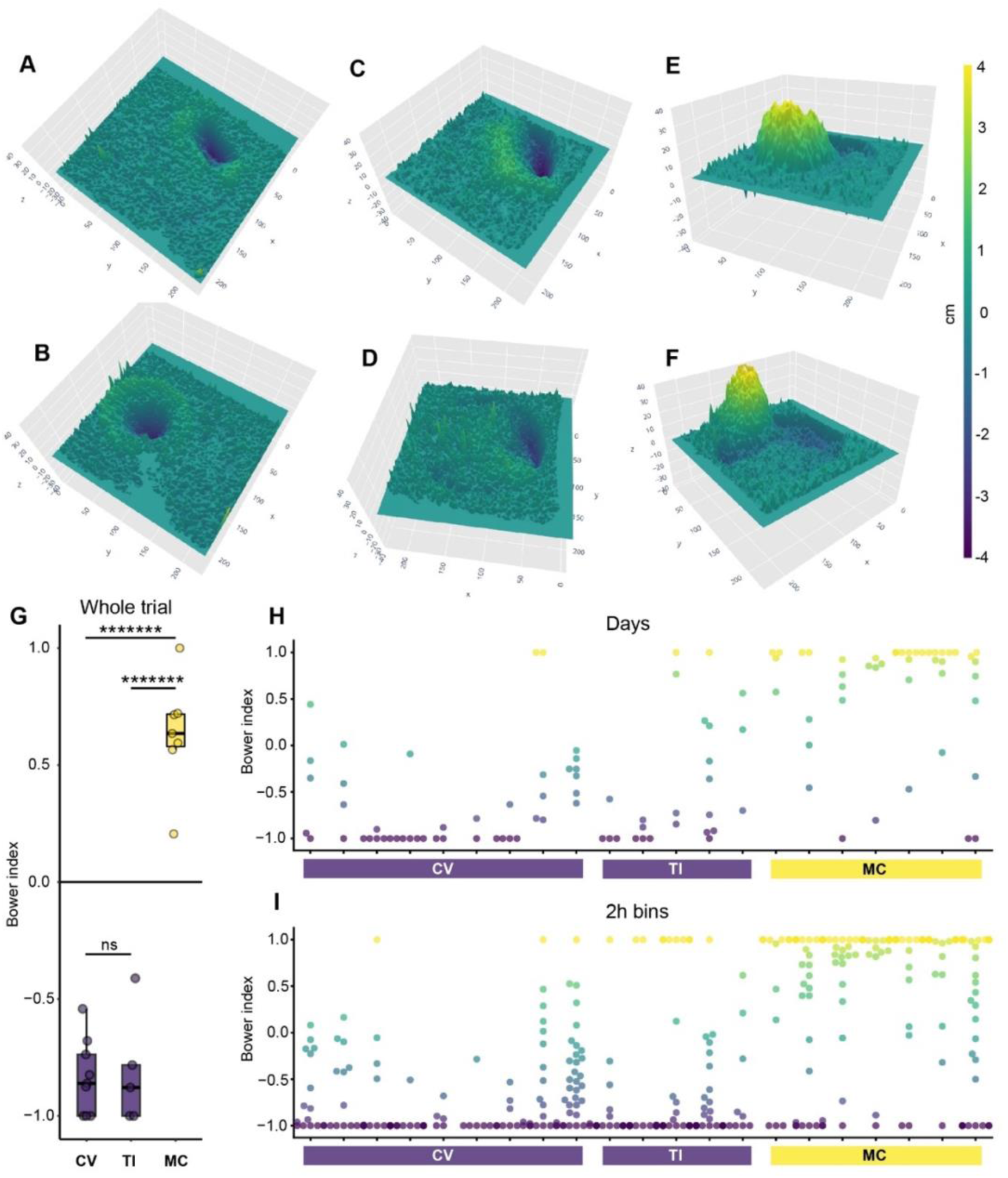
Depth sensing reveals natural species differences in bower construction at multiple timescales. Depth sensing allows visualization and analysis of bower construction across multiple species and timescales. Representative 3D reconstructions show pits constructed by *Copadichromis virginalis* (CV, n=9; A,B), pits constructed by *Tramitichromis intermedius* (TI, n=5; C,D), and castles by *Mchenga conophoros* (MC, n=7; E,F; all z-axes amplified for visual effect). The Bower Index, a measure of the directional bias in bower construction, revealed strong species differences in the final bower structures (whole trial change, e.g. A-F). MC exhibited a positive bias (extreme depth change regions tended to be elevated) and differed strongly from both TI and CV, which exhibited negative biases and did not differ significantly from each other (G). Strong species differences in structural development were also present when depth data was analyzed at daily (H) and hourly (I, 2-hour bins) timescales, mirroring the direction of differences observed for whole trial change. Each point represents the observed Bower Index for a single time bin for within one trial. Different individuals are separated into columns along the x-axis, grouped by species. The color of each point reflects the Bower Index for that time bin (−1, purple, pure pit-like depth change; 1, yellow, pure castle-like depth change).

#### 3.2.2 Biological validation

*Depth sensing captures natural species differences in bower structures* We next tested whether our depth sensing system could detect natural species differences in bower structures. To do this, we compared depth change in bower trials among three species: two pit-digging species (*Copadichromis virginalis*, n=9; *Tramitichromis intermedius*, n=5) and one castle-building species (*Mchenga conophoros*, n=7). We calculated a “Bower Index” to analyze the final bower structure in each trial (**Figure 3G**). Briefly, the Bower Index is a ratio of the net depth change in above threshold regions (change can be positive and negative) to the total volume change in above threshold regions (all change is considered positive). The Bower Index is thus a measure of directional (elevation vs. depression) bias in above threshold regions. This analysis revealed strong species differences in bower structures (One-way ANOVA, p=5.42×10^−11^), with all pit-digging individuals exhibiting a negative Bower Index (14/14), and all castle-building individuals exhibiting a positive Bower Index (7/7). Post-hoc Tukey’s HSD tests revealed that pit-digging species did not differ significantly from each other (CV vs. TI, Tukey’s HSD, p=0.98; Fig. 4G), but the castle-builder *Mchenga conophoros* differed strongly from both pit-digging species (MC vs. TI, Tukey’s HSD, p=1.68×10^−9^; MC vs. CV, Tukey’s HSD, p=1.16×10^−10^). Strong species differences in structural development were also present when depth data was analyzed at daily (24-hour bins; One-way ANOVA, p=4.95×10^−8^; H) and hourly (2-hour bins; One-way ANOVA, p=1.62×10^−11^; I) timescales, mirroring the same pattern of differences among species that was observed at the whole trial level (**Figure 3H,I**).

**Figure 4.**
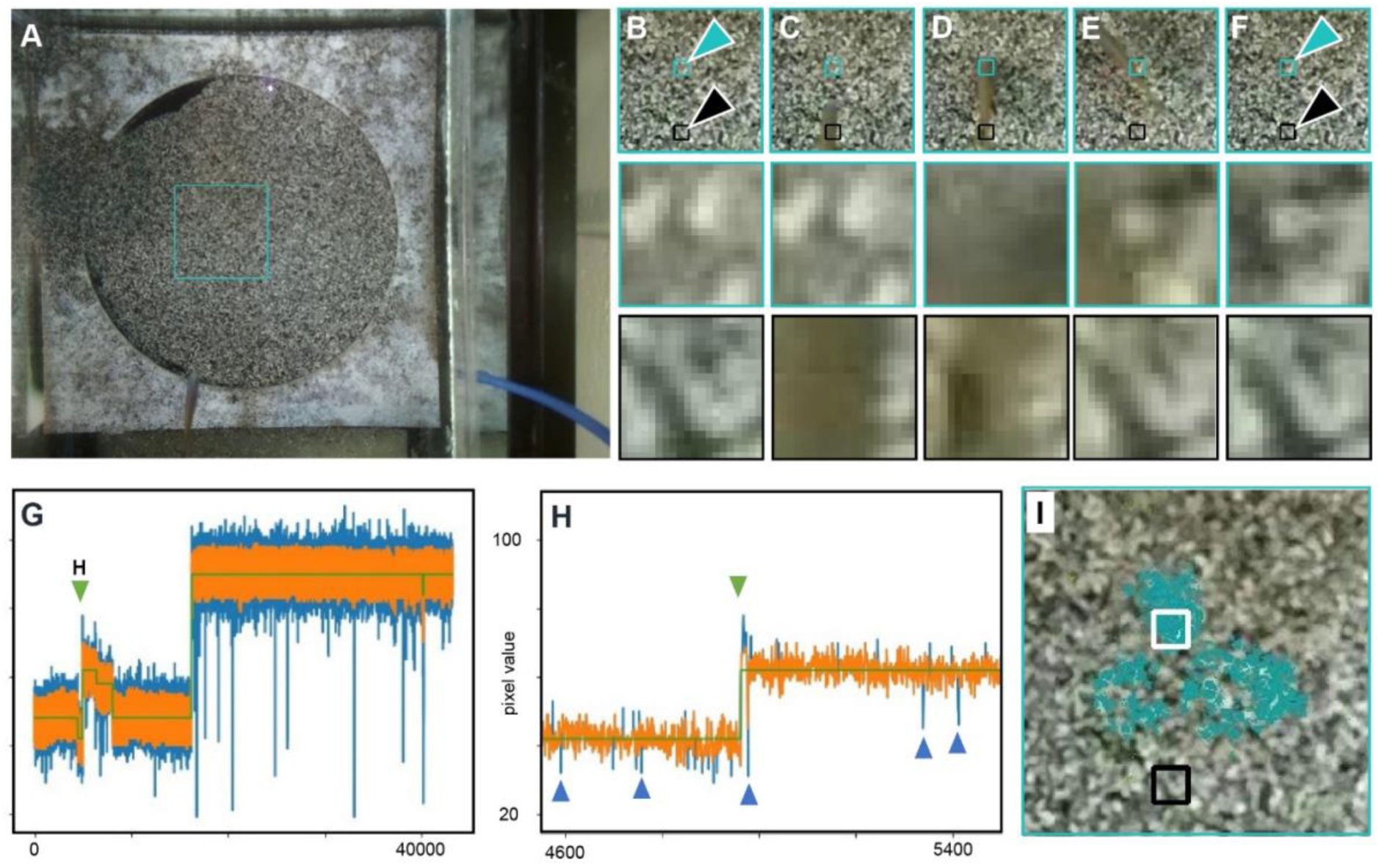
Automated detection of sand change from video data. The sand in this behavioral paradigm is composed of black and white grains (e.g. as seen in panel A), and therefore sand manipulation events during bower construction cause permanent rearrangement of the black and white grains at specific locations. We aimed to detect these events by processing whole video frames (A, with turquoise box indicating an example region of interest) sampled once per second, and tracking the values of individual pixels throughout whole trials. Fish swimming over sand cause transient changes in pixel values (e.g. B-F, black arrows indicate an example location of a fish swimming over the sand; the bottom row depicts a zoomed in 20×20 pixel view of a location that the fish swims over, sampled from representative frames across four seconds). In contrast, sand manipulation behaviors cause enduring changes in pixel values (e.g. B-F, turquoise arrow indicate an example location of a fish scooping sand; the middle row contains a zoomed in 20×20 pixel view of a location where the fish scoops sand). We used a custom Hidden Markov Model to identify all enduring state changes for each pixel throughout entire videos (G, green line indicates HMM-predicted state, orange line indicates natural variance in pixel value, and blue lines indicate transient fluctuations beyond the pixel’s typical range of values likely caused by fish swimming or shadows). Because fish swim over the sand frequently, a large number of transient changes are ignored (e.g. pixel value fluctuations indicated by blue arrows in panel H), while enduring changes are identified (e.g. pixel value change indicated by green arrow in Panel H). Density-based clustering identifies high-density spatiotemporal clusters of HMM+ pixels, or putative sand change events, for further analysis (turquoise pixels represent one sand change cluster, I; region of interest and boxes correspond to A-F).

### 3.3 Video data

#### 3.3.1 System validation

##### Automated identification of sand change from video data

To investigate bower construction on more acute time scales, we created tools to track sand change events from video data across whole trials. We took advantage of the multi-color sand (composed of black and white grains) in our setup: each time a fish contacts the sand it causes an enduring spatial rearrangement of the black and white grains of sand, changing the corresponding pixel color value from top-down video. In contrast, a fish swimming over the sand (and any shadows it casts) only causes transient changes in pixel values (**Figure 4A-F**), after which each pixel returns to its original value (i.e. the same value before the fish swam by). We found that a Hidden Markov Model (HMM) could identify enduring sand rearrangements while simultaneously ignoring transient changes caused by swimming fish (**Figure 4G,H** and **S5**), and further that groups of spatially and temporally concentrated “clusters” of sand change pixels could be identified using density-based clustering (**Figure 4I, S6, S7**, and **S8**). This approach allowed us to map the times and spatial locations of thousands of fish-mediated sand manipulations on each day of each bower trial. Manual review confirmed that the vast majority of predicted sand change clusters (>90%, 13,288/14,234 analyzed events) were true sand change events caused by fish behaviors, with the remaining portion including reflections of events in the glass, shadows caused by stationary or slow-moving fish, or in rare cases small bits of food, feces, or other debris settling on the sand surface.

##### Automatic classification of cichlid behaviors with action recognition

Because bowers are constructed through thousands of spatial decisions over many days, manually scoring full trials would be impractically labor intensive, and we therefore aimed to automatically identify bower construction behaviors from video data. However, scooping and spitting sand during bower construction represents only a subset of behaviors that cause sand change in our paradigm. For example, feeding behaviors are performed by both males and females and also involve scooping and spitting sand, and are expressed frequently throughout trials. Quivering and spawning behaviors, in which a male rapidly circles and displays for a gravid female, are less frequent but also cause large amounts of sand change. We therefore aimed to automatically separate bower construction events from other behaviors that cause sand change. We first evaluated several methods for distinguishing bower scoops and spits from each other and from other types of events, including analysis of spatial properties of sand change clusters (e.g. cluster size, see **Figure S10**), and feature extraction from short video clips generated for events. While these methods revealed differences between behavioral categories, our preliminary analyses suggested they were insufficient for accurately classifying behaviors. We then turned to a deep learning approach and assessed whether 3D ResNets, which have been recently shown to accurately classify human actions from video data (Qiu, Yao et al. 2017), could accurately distinguish fish behaviors that cause sand change in our paradigm.

To create a training set for the 3D ResNet, we generated short cropped video clips centered spatially and temporally around each sand change event from a subset of seven behavioral trials, representing seven individuals, three species, and one pit-castle hybrid cross. A trained observer manually annotated a randomly sampled subset of 14,234 video clips (∼2,000 per trial). Each clip was classified into one of ten categories (bower scoop, bower spit, bower multiple, feed scoop, feed spit, feed multiple, drop sand, quivering, fish other, and other; for operating definitions used for all behaviors see Supplementary Materials subsection “Behavioral definitions”). Feeding was the most frequently observed behavior, accounting for nearly half of all clips (46.9%, 6,672/14,234 annotated clips; feeding scoops, 15.2%; feeding spits, 11.5%; multiple feeding events, 20.2%). Bower construction behaviors were the next most prevalent (19.5%; bower scoops, 9.4%; bower spits, 8.1%; multiple bower construction events, 1.9%). Quivering and spawning events were the least frequently observed, accounting for just 2.6% of all clips. The remainder of sand change events were annotated as either sand dropping behavior (5.6%), “other” behaviors (e.g. brushing the sand surface with the fins or the body; 18.8%), or shadows/reflections (6.6%).

A 3D ResNet was then trained on 80% (∼11,200 clips) of the data, and the remaining 20% of the data was used for testing (∼2,800 clips). To place the ResNet predictions in the context of human performance, we also measured the accuracy of a previously naive human observer that underwent 12 hours of training and then manually annotated a test set of 3,052 clips from three trials and all ten behavior categories. The 3D ResNet achieved ∼77% accuracy on the test set, which was comparable to a newly trained human observer (∼80% accuracy, 2,456/3,052 clips). Confidence for 3D ResNet predictions on the test set ranged from 22.1-100%, and confidence tended to be greater for correct predictions (mean confidence 92.93±0.279%) than for incorrect predictions (mean confidence 78.28±0.074%) (**Figure S11**). We found an imbalance in the distribution of incorrect predictions across categories (**Figure 5A**). For some categories, such as “build multiple”, “feed multiple”, and “fish other”, video clips could contain behaviors that also fit into other categories. For example, a “feed multiple” clip by definition contains multiple feeding scoop and/or feeding spit events, a “bower multiple” clip contains multiple bower scoops and/or bower spits, and a “fish other” clip may contain a bower scoop and a fin swipe (or some other combination of behaviors). We found that erroneous “within building” category predictions for build multiple, “within feeding” predictions for feed multiple, and “fish other” predictions accounted for ∼82% of all incorrect predictions. We further found that setting a confidence threshold of 90% excluded most (∼62%) incorrect predictions but included most (70%) correct predictions, including ∼86% of correct bower scoop predictions and ∼88% of correct bower spit predictions. 69% of all predictions were above the 90% confidence threshold, and overall accuracy for these high-confidence predictions was ∼87% (**Figure S11**).

**Figure 5.**
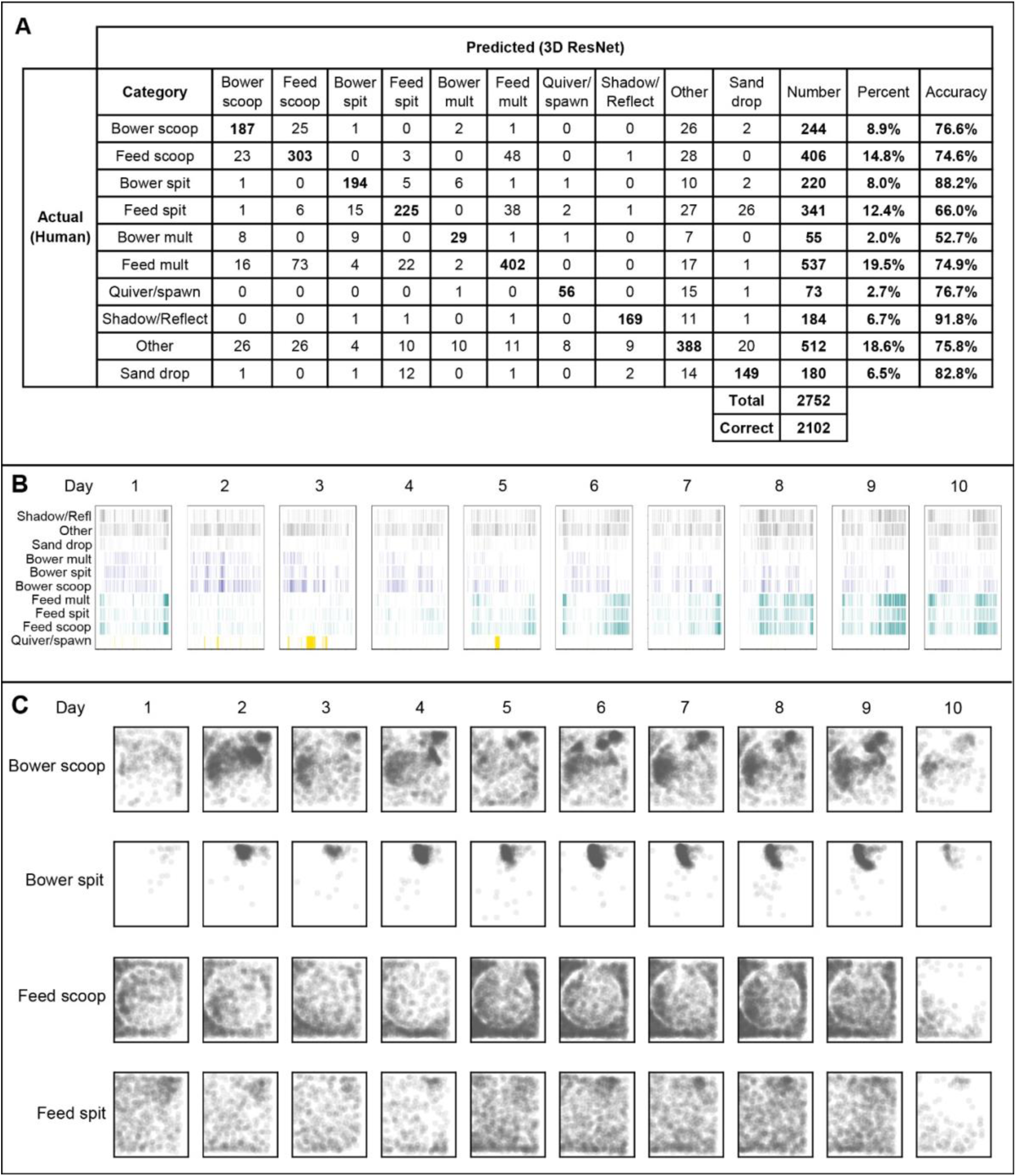
Deep learning and prediction accuracy of cichlid behaviors. A confusion matrix for predictions on the test dataset shows that predictions made by the 3D ResNet tended to match human annotations across all ten behavioral categories (A, emboldened diagonal values indicate the number of agreements between human annotations and 3D ResNet predictions for each category). By applying the 3D ResNet across full trials, bower construction, feeding, and spawning behaviors can be temporally mapped over long timescales, spanning >100 hours of video data (B). Spatially mapping 3D ResNet-predicted bower construction and feeding behaviors reveals distinct spatial distributions among behaviors that are often indistinguishable to untrained human observers (C).

##### Spatial and temporal mapping of behavioral events

Because all behavioral predictions were linked to individual sand change clusters, each event was associated with a unique timestamp and pixel coordinate location within video data. Temporally mapping behavioral events revealed that behaviors were expressed non-uniformly in time (**Figure 5B**). Similarly, spatially mapping behavioral events revealed distinct patterns for each category, including strikingly different spatial patterns between construction behaviors versus feeding behaviors (**Figure 5C**).

### 3.4 Combined video and depth data

#### 3.4.1 System validation

##### Registration links behavioral events to depth data through time

We next spatially and temporally aligned video and depth data for the same seven trials used to train the CNN. We used RGB images collected with the Kinect for spatial registration of video and depth data, and we used time stamps assigned by the Raspberry Pi for temporal alignment. We found that most (∼56%) of CNN-predicted events could be linked to sand surface height at the corresponding time and location. We also found a large proportion (∼44%) of events could not be linked to surface height, which was not surprising because the video FOV included the glass walls outside the sand tray, and ∼10% of the sand surface was not captured by the Kinect. We observed a bias in the types of events that could not be linked to depth change values, with just five categories accounting for ∼87% of these predictions (shadow/reflections: 31.9%, other: 21.3%, feed multiple: 14.2%, feed scoop: 12.7%, and feed spit: 6.4%). This was not surprising, as fish frequently feed along the periphery against the glass walls, and “shadow/reflections” includes reflections of events in the glass. In contrast, bower scoops and spits represented a small minority of these events (bower scoop: 3.4%, bower spit: 4.1%), supporting high quality depth data in regions where the males constructed bowers.

#### 3.4.2 Biological validation

##### Agreement between action recognition and depth sensing

Using registered video and depth data, we tested how 3D ResNet-predicted scoop and spit events mapped onto bower structures identified from depth data (**Figure 6**). Because pits are excavated by scooping sand, we predicted that a greater number of scoops compared to spits would occur within the most extreme depth change regions of interest (bower ROIs) in pit-diggers, and that the opposite pattern would be observed in castle-builders. To test this, we compared the number of scoops and spits observed inside and outside the bower ROI for each of the five parental trials analyzed by the 3D ResNet (n=1 CV, n=2 TI, n=2 MC). Indeed, in pit-diggers we observed ∼15x more CNN-predicted scoops versus spits within daily bower ROIs and this pattern was highly significant within each subject (CV: 273 scoops vs. 20 spits, χ^2^=311.35, p<2.2×10^−16^; TI subject 1: 339 scoops vs. 16 spits, χ^2^=127.2, p<2.2×10^−16^; TI subject 2: 602 scoops vs. 60 spits, χ^2^=377.28, p<2.2×10^−16^). This pattern was flipped in castle-builders, with ∼5.5x more CNN-predicted spits versus scoops occurring within daily bower ROIs (MC subject 1: 242 scoops vs. 2,208 spits, χ^2^=5554.2, p<2.2×10^−16^; MC subject 2: 260 scoops vs. 462 spits, χ^2^=208.92, p<2.2×10^−16^).

**Figure 6.**
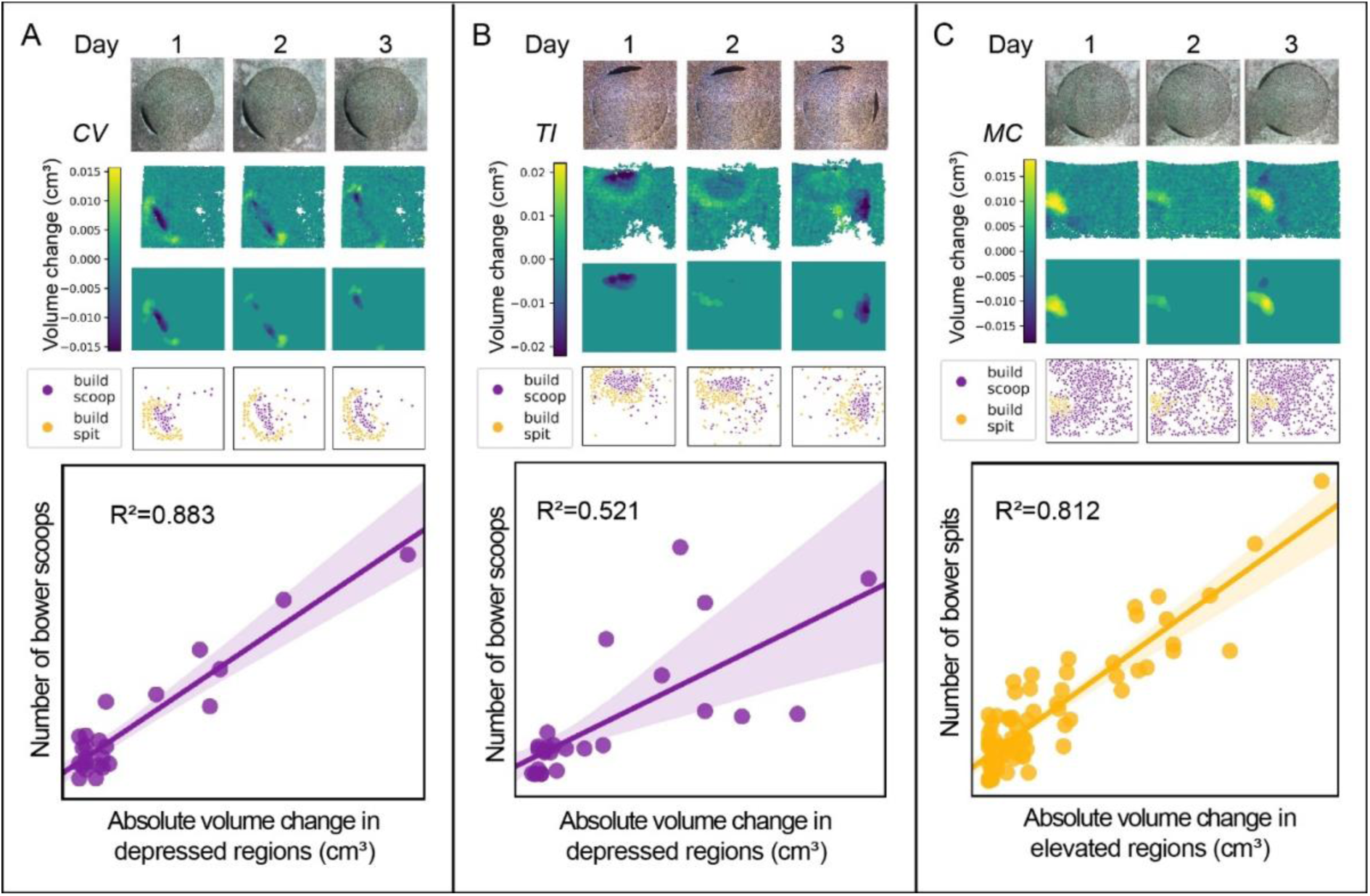
CNN-predicted behavioral events predict bower structures. RGB images collected with the Kinect (first row, A-C) were registered to RGB frames collected with the Raspberry Pi Camera to spatially align video and depth data. Daily depth change data (second row, A-C) was analyzed to identify above-threshold regions (third row, A-C). In pit-diggers (A, B), a greater proportion of CNN-predicted bower scoops versus bower spits mapped onto extreme height change regions (overlap of third and fourth rows), whereas in castle-builders the reverse was true: a greater proportion of bower spits versus bower scoops mapped onto extreme height change regions. In pit-diggers, the number of bower scoops per hour was strongly and positively correlated with the total volume change in that hour (e.g. see representative regression plots for individual trials in A, B), whereas in castle-builders the number of bower spits per hour was strongly and positively correlated with the total volume change in that hour (regression plot, C).

We also investigated whether the temporal distribution of CNN-predicted events was associated with the temporal development of the bower structure. In pit-diggers, we found that the number of hourly bower scoops was strongly and positively correlated with the hourly volume change in depressed regions (R^2^=0.597, p<0.00001); whereas in castle-builders, the number of hourly bower spits was strongly and positively correlated with the hourly volume change in elevated regions (R^2^=0.690, p<0.00001, representative trials shown in **Figure 6**). Taken together, these data demonstrate agreement between two orthogonal data streams, and show that behaviors identified through action recognition are predictive of the spatial, geometric, and temporal development of the bower structure measured through depth sensing.

### 3.5 New biological insights

#### Bower behaviors are spatially repeatable

We used depth data to test a new biological dimension of bower building behavior: do males construct their bowers in the same spatial location across trials? To do this, we tracked seven subject males across multiple trials, between which the male was temporarily removed, the bower was abolished, the sand surface was smoothed, and the male was reintroduced (e.g. see **Figure 7A-E**, first and second columns representing first and second trials, respectively). For each repeatability subject, we calculated the spatial overlap of above-threshold regions between trials (e.g. see **Figure 7A-E**, third column). We found that the observed spatial overlap between trials was significantly greater than the overlap expected by chance (**Figure 7F**; 23.4±4.30% overlap between repeatability trials versus 2.5±0.64% overlap expected by chance; paired t-test, p=0.000264; n=14, pooled by species/cross). The direction of this effect was the same within each species and each cross (Supplementary Table 1). Despite small sample sizes, this effect was also significant within CV alone as revealed by a paired t-test (n=5 pairs of repeatability trials, p=0.0228).

**Figure 7.**
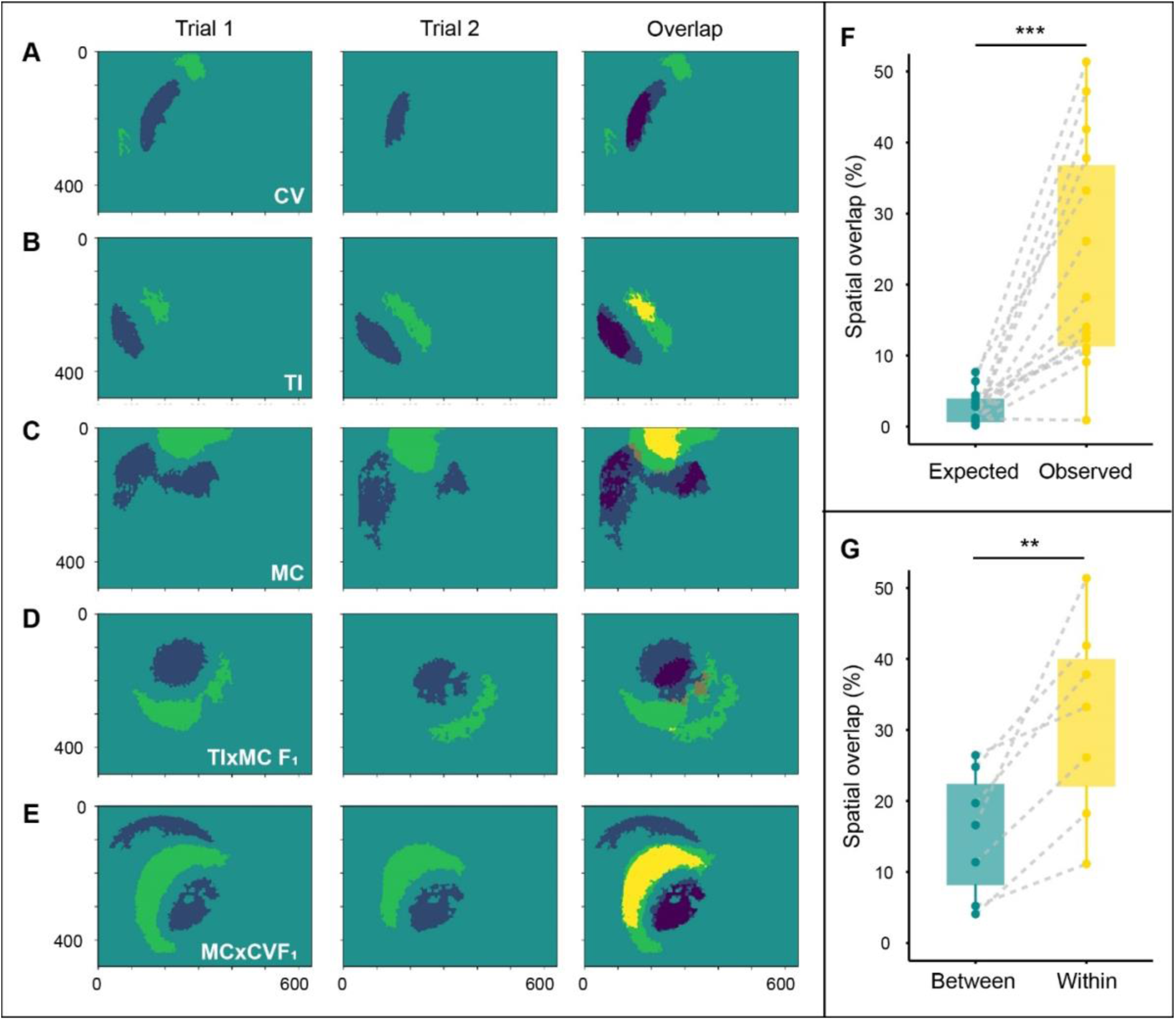
Bower construction is spatially repeatable. Bower construction behaviors are spatially repeatable. Analyzed males represented three species (top-down depth sensing data; *Copadichromis virginalis*, n=5, A; *Tramitichromis intermedius*, n=2, B; *Mchenga conophoros*, n=4, C) and two pit-castle F_1_ hybrid crosses (TIxMC F_1_, n=2, D; MCxCV F_1_, n=1, E). Following Trial 1 (A-E, first column), males were temporarily removed, the sand tray was reset, and males were reintroduced to the same tank for repeatability trials (A-E, Trial 2, second column). Spatial overlap was calculated as the ratio of shared above-threshold (A-E third column, bright yellow) and below threshold (A-E third column, dark blue) regions, relative to the total above and below threshold regions in either trial. Spatial overlap of above-threshold regions between trials was significantly greater than overlap expected between randomly distributed regions of the same size (F). Overlap between trials for individual males was greater than overlap between trials for different males of the same species tested in the same tank (G).

Spatially repeatable bower construction could be driven by a spatial memory of the bower location maintained across trials, or by tank-specific factors that might cause some locations within each tank to be generally more preferable for bower construction. To investigate these two possibilities, we compared pairs of repeatability trials with pairs of trials in which different subjects of the same species were tested in the same tank. First, we found that overlap between different males of the same species tested in the same tank was greater than expected by chance (10.3±3.04% spatial overlap observed versus 3.4±1.67% expected by chance), supporting that some locations within each tank were generally more preferable for bower construction, across subjects. However, in 7/7 cases, we found that spatial repeatability was also stronger within subjects than between species-matched subjects tested in the same tank (**Figure 7G**; p=0.0045; pooled by species/cross: CV, n=3; MC, n=1; TIxMCF_1_, n=2; MCxCVF_1_, n=1), consistent with the idea that spatial memory also plays a role in bower (re)construction. Despite small sample sizes, this effect was also significant within CV only (n=3, paired t-test, p=0.0035).

#### Bowers structures develop non-uniformly in space

The spatial repeatability of bower construction raises the question as to what rules guide male decision-making before a structure is present, and later after a visually salient structure has begun to develop. One possibility is that males construct bowers in a spatially uniform manner over the full course of construction—within each punctuated burst of activity, the bower structure develops proportionately and spatially uniformly toward its final form. A second possibility is that the bower arises in a spatially non-uniform manner, with different regions of the bower developing disproportionately relative to one another.

To investigate these models, we developed a Spatial Uniformity Index (SUI) to measure the disparity between the actual structural change observed on each day of bower construction, and the structural change expected under the assumption of perfect spatial uniformity (1=perfectly uniform, 0=zero spatial uniformity) based on the daily volume of sand moved. In other words, if 20% of total volume change occurs on the first day of construction, and the final bower structure develops to 20% of its final height, then the SUI for the first day will equal 1. Analysis of the SUI across all above-threshold days for all bower trials (n=29 total; CV, n=9; TI, n=5; MC, n=7; MCxCV F_1_, n=3; TIxMC F_1_, n=5) provided two new insights into bower construction (**Figure 8**). First, linear mixed-effects regression with SUI as the outcome variable; species, day, and the interaction between species and day as fixed effects; and subject as a random effect revealed that the SUI was much closer to 0 than 1 in all three species (regression estimate of mean for CV=0.19±0.036, TI=0.12±0.051, MC=0.12±0.037) and both hybrid crosses (regression estimate of mean for (Satterthwaite’s method, F=5.47, Tukey’s p=0.00057), but not of species (Satterthwaite’s method, F=0.74, Tukey’s p=0.57), or the interaction between species and day (Satterthwaite’s method, F=0.72, Tukey’s p=0.77) on spatial uniformity. Spatial uniformity was especially low on the first day of bower construction (regression estimate for Day 1, 0.082±0.0300) and was nearly three times more uniform on the second day (regression estimate for Day 2, 0.24±0.0314), a shift that gradually tapered off (Day 3: 0.20±0.0346; Day 4: 0.15±0.0350; Day 5: 0.11±0.0487). Post-hoc analysis of pairwise differences among days revealed the increase in spatial uniformity from Day 1 to Day 2 to be significant (t=-4.313, Tukey’s p=0.0004), and from Day 1 to Day 3 to be significant (t=-2.938, Tukey’s p=0.034), but no other pairwise differences between days were significant. Although our calculation accounted for differences in volume change from day to day, we were still concerned that the shift in SUI could be an unexpected byproduct of less sand being moved on the first day of bower construction compared to other days. To directly test this, we added daily volume change directly as a fixed effect to the same model. This model showed that daily volume of sand moved was not associated with SUI (Satterthwaite’s method, F=0.12, Tukey’s p=0.73), and that SUI was strongly associated with day even when directly controlling for daily volume change in the model (Satterthwaite’s method, F=5.42, Tukey’s p=0.00075). Taken together, these data support a significant shift in spatial decision-making patterns during the early stages of bower construction, perhaps corresponding to the transition from constructing on a flat sand surface to constructing when a structure is present.

**Figure 8.**
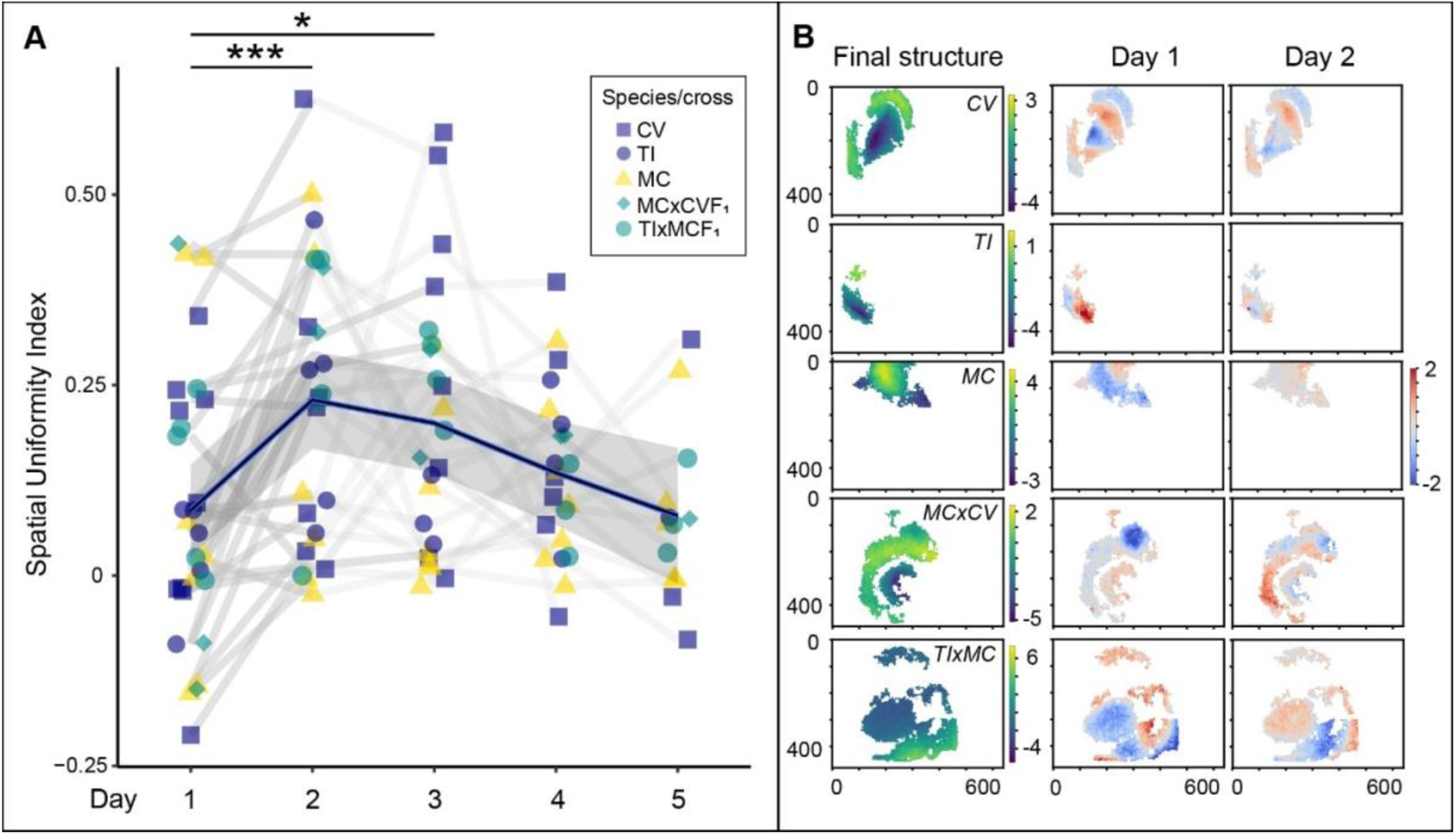
Spatial patterns shift over the course of bower construction. Analysis of spatial uniformity revealed an overall trend of low uniformity (closer to 0 than 1, A) in all species and hybrid crosses over the course of bower construction. Uniformity was lowest on the first day of bower construction and significantly increased by nearly threefold on the second day (A; Tukey’s p=0.018), before gradually tapering off by the fifth day (A; linear mixed-effects mean and standard error estimates indicated by black line and gray band, respectively). Shifts in uniformity from Day 1 to Day 2 are shown for representative subjects from each species and hybrid cross (Analysis of top-down depth sensing data, B). The left column shows the final structure in above threshold regions for each trial. The second and third columns show spatial patterns of non-uniformity on Days 1 and 2, respectively, or the disparity between the actual structural change and the structural change expected under the assumption of perfect spatial uniformity (red indicates regions in which height increased more than expected, blue indicates regions in which height decreased more than expected). Units for all heatmaps are cm, and pixels are marked on the x and y axes of plots in (B).

#### Strong shifts in behavioral and social contexts across full trials

We also investigated behavioral and social contexts across whole trials (**Figure 9A**). First, we investigated whether different behaviors covaried strongly with one another through time. This analysis showed a clear pattern of distinct behavioral contexts through time, driven by three behavioral clusters. One cluster was driven by strong covariation among feeding behaviors and sand dropping behavior, a second cluster was driven by strong covariation among bower construction behaviors and “other” behaviors, and spawning/quivering behaviors covaried weakly with both feeding and bower construction behaviors. Taken together, these data support that feeding, bower construction, and mating contexts occur distinctly through time. Pairwise Pearson’s R values and corresponding p-values are shown in **Supplementary Table 2**.

**Figure 9.**
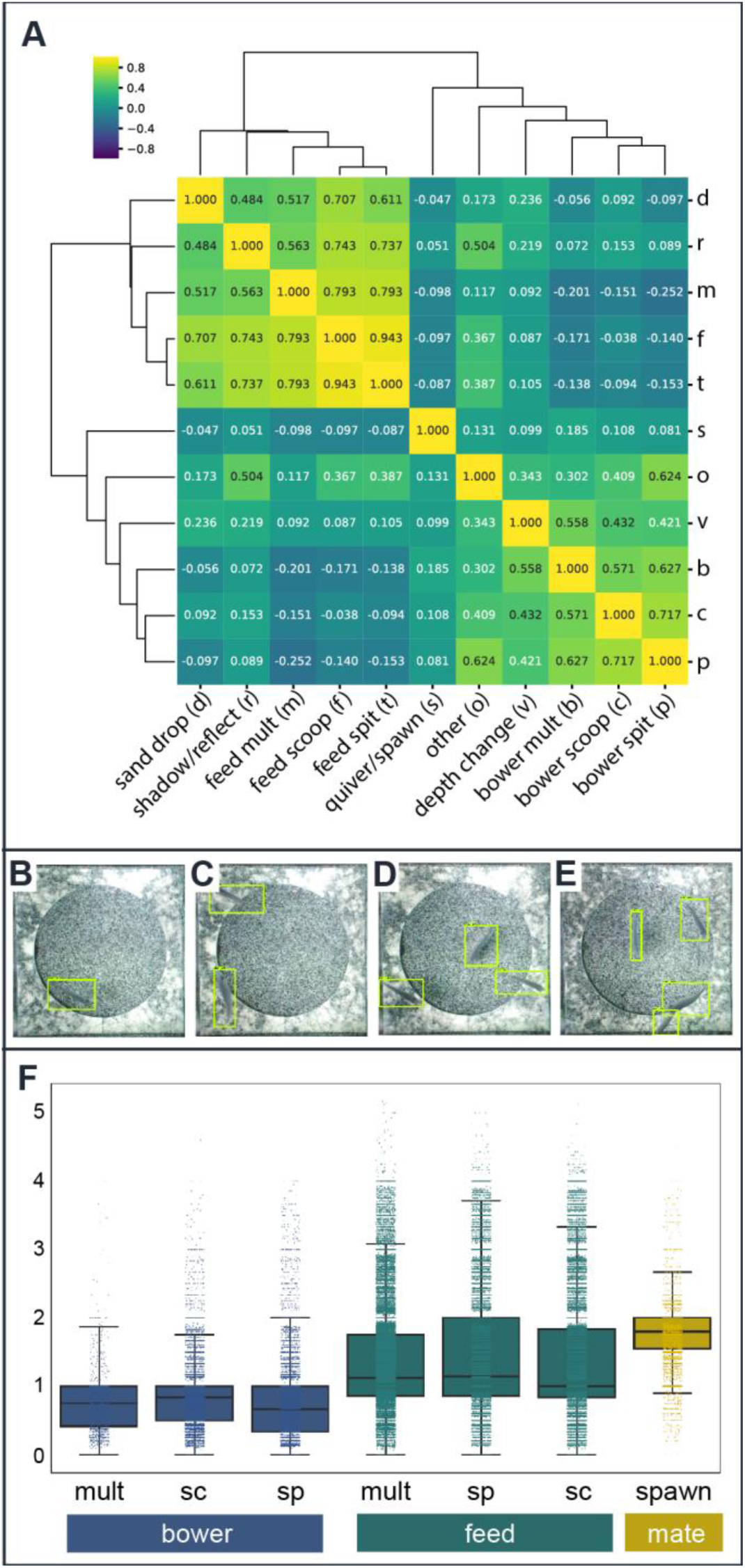
Distinct behavioral and social contexts across whole trials. Patterns of covariation among 3D ResNet-predicted behavioral events support strong shifts among three behavioral contexts across whole trials, corresponding to feeding, bower construction, and spawning behaviors (A, Pearson’s R values shown for each pairwise correlation). A Faster R-CNN detected and counted fish above the sand tray from whole video frames, with green outlines indicated predicted fish (B-E; 1, 2, 3, and 4 fish detected, respectively). Analysis of the number of fish present above the sand tray during 3D ResNet-predicted behavioral events revealed strong differences in fish count across behavioral categories (E;p<2×10^−16^). Fish counts were lowest during bower construction, greater during feeding, and greatest during spawning, supporting dynamic and intertwined behavioral and social contexts across whole trials.

We next used object recognition to count fish in order to test whether the social dynamics among males and females differed between these behavioral contexts. To do this, we trained Faster-RCNN networks to identify and count fish using ∼1800 manually annotated frames with an accuracy of ∼95% (**Figure 9B**). Linear mix-effects regression with fish count as the outcome variable, behavior as a fixed effect, and day nested within individual nested within species as a random effect, revealed that the number of fish present above the sand tray differed strongly across behavioral contexts (F=2285.9, p<2.2×10^−16^) (**Figure 9C**). The fewest fish were present during bower behaviors (average fish count regression estimates for build scoop=0.99±0.118, build spit=0.88±0.118, build multiple=0.87±0.118); a greater number tended to be present during feeding behaviors (feed scoop=1.24±0.118, feed spit=1.27±0.118, feed multiple=1.21±0.118); and the greatest number of fish, on average, were present during spawning behaviors (1.93±0.118). Post-hoc pairwise comparisons revealed significant differences between all behaviors with the exception of bower spit versus bower multiple events (t=-0.396, p=0.997). Taken together, these data support strong shifts in behavioral and social contexts across full trials, driven by distinct periods of feeding, bower construction, and spawning.

#### Sequential expression of parental behaviors in pit-castle F_1_ hybrids

We also intercrossed pit and castle species and investigated expression of parental behaviors in pit-castle F_1_ hybrid offspring (**Figure 10**). We have previously observed in one pit-castle F_1_ hybrid cross (*Mchenga conophoros* dam *x Copadichromis virginalis* sire, MCxCV) that males appear to express both parental behaviors in sequence, first digging a pit and then building a castle; however this transition has not previously been quantified. We aimed to measure this transition in two “reciprocal” pit-castle hybrid crosses: MCxCV, and a second cross, *Tramitichromis intermedius* dam x *Copadichromis virginalis* sire (TIxMC), which has not been previously recorded (MCxCV, n=3; and *Tramitichromis intermedius* dam x *Mchenga conophoros* sire, n=5). Analysis of the Bower Index from day to day revealed a trajectory in which F_1_ hybrid males transitioned from a pit-like Bower Index on Day 1 (MCxCV, -0.85±0.075; TIxMC, n=5, -0.58±0.194) to a castle-like Bower Index by Day 5 (MCxCV, 0.44±0.155; TIxMC, n=5, 0.40±0.289). A linear mixed-effects model with Bower Index as the outcome variable; cross, day, and the interaction between cross and day as fixed effects; and subject as a random effect, revealed a strong effect of day (F=14.30, p=2.8×10^−6^) but not of cross (F=0.024, p=0.88) or the interaction between cross and day (F=1.11, p=0.37) on the Bower Index. Post-hoc analysis showed that the transition from Day 1 to Day 5 was significant in both crosses (Day 1 vs. Day 5: MCxCV, t=-4.44, Tukey’s p=0.0038; TIxMC, t=-4.565, Tukey’s p=0.0041). Taken together, these data show a strong and similar transition from pit-biased to castle-biased behavior in both F_1_ crosses.

**Figure 10.**
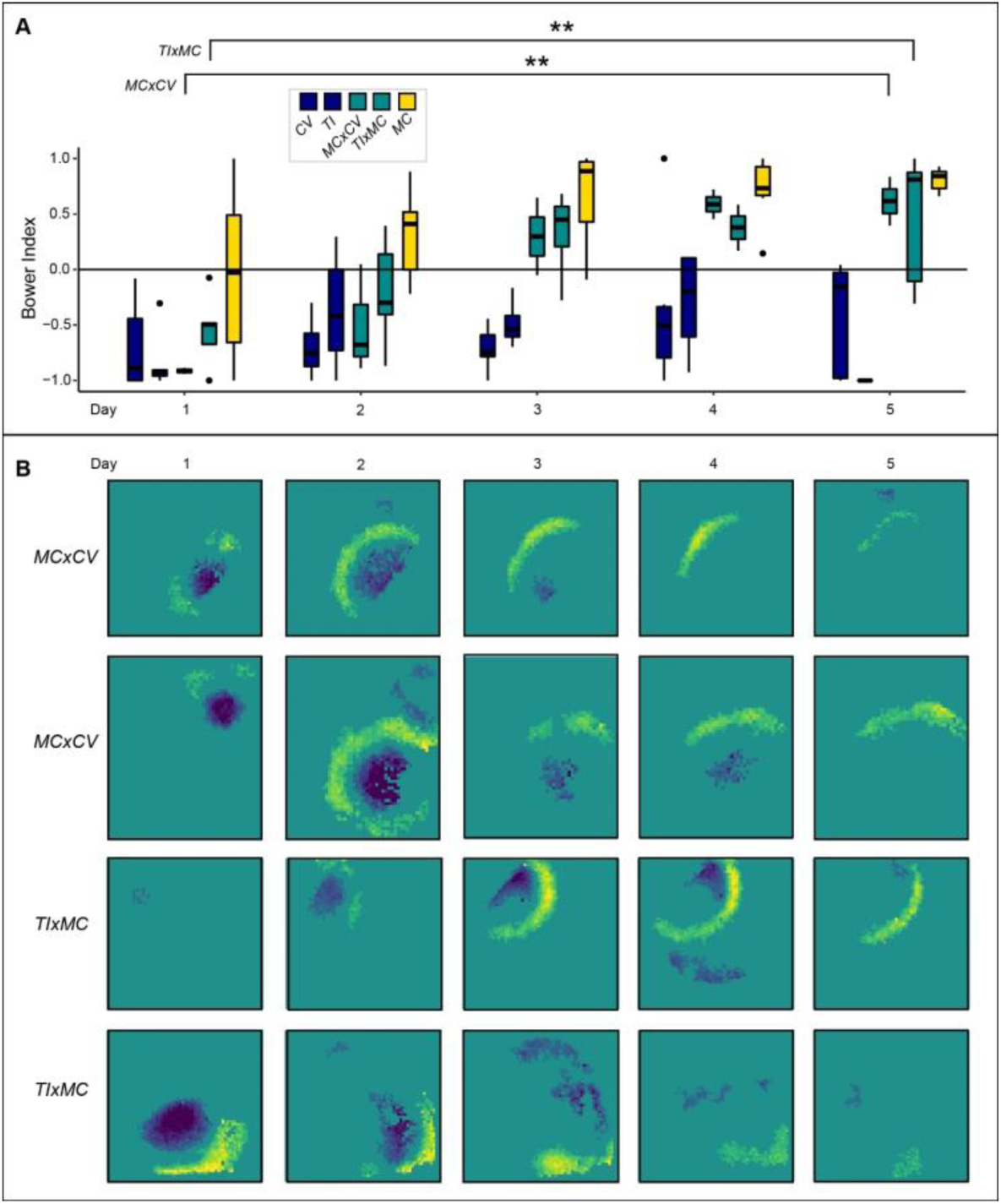
Interspecies pit-castle F_1_ hybrid males sequentially express pit-digging then castle-building. Analysis of the Daily Bower Index through time shows that pit- castle F_1_ hybrids (turquoise boxes) transition from a pit-like behavioral phenotype on Day 1 to a castle-like behavioral phenotype on Day 5 (A; the first dark blue box represents CV on each day, the second dark blue box represents TI, the first turquoise box represents MCxCV F_1_ hybrids, the second turquoise box represents TIxMC F_1_ hybrids, and the yellow box represents MC). The Bower Index significantly increased from Day 1 to Day 5 in both F_1_ crosses (MCxCV, p=0.0038; TIxMC, p=0.0041). Plots of above-threshold depth change illustrate the development of pit-like regions on Day 1, with a gradual shift to more castle-like development by Day 5 (B, first column; each row represents a different trial and F_1_ subject).

To place this transition in the context of parental behavior, we performed simple oneway t-tests to assess whether the F_1_ Bower Index was greater compared to pit-diggers, or less compared to castle-builders across days. Because we found no evidence for any behavioral difference between crosses, F_1_ subjects were pooled for comparison with parental species. The Day 1 Bower Index in pit-castle F_1_ hybrids (n=8) did not differ significantly from the Day 1 Bower Index in either pit-digging species (vs. CV, n=9, p=0.40; vs. TI, n=5, p=0.19), but was significantly less than castle-builders (vs. MC, n=7, p=0.038). By Day 2, the F_1_ bower index was greater than pit-digging CV (p=0.038), indistinguishable from pit-digging TI (p=0.43), and still significantly less than castle-building MC (p=0.012). By Day 3 the pattern had fully reversed, with the F_1_ Bower Index significantly greater than both pit-digging species (vs. CV, p=3.7×10^−5^; vs. TI, p=0.0033) but no longer distinguishable from castle-builders (vs. MC, p=0.097), and this pattern persisted through Day 5. Taken together, these data support sequenced expression of parentally-biased behaviors in F_1_ hybrid males.

## Discussion

Construction behaviors are excellent natural models of long-term goal-directed decision-making in dynamic environments, but it is difficult to simultaneously measure a developing structure and an animal’s behavioral decisions over long timescales. However, new tools are providing entry points for automated measurements of natural behaviors in the lab. For example, static poses and positions of animals are being tracked through time in increasingly complex environments (Dell, Bender et al. 2014, Robie, Seagraves et al. 2017, Hughey, Hein et al. 2018). Depth sensing, radio-frequency identification (RFID) tagging, and additional cameras have been used in conjunction with standard video data to track animals in complex social environments in which occlusions regularly occur (Ardekani, Biyani et al. 2013, Weissbrod, Shapiro et al. 2013, Hong, Kennedy et al. 2015, Macfarlane, Howland et al. 2015, Wiltschko, Johnson et al. 2015). Software tools such as DeepLabCut and idTracker.ai also enable pose estimation and positional tracking from video data in which animals are behaving in complex environments (Perez-Escudero, Vicente-Page et al. 2014, Mathis, Mamidanna et al. 2018, Nath, Mathis et al. 2019, Romero-Ferrero, Bergomi et al. 2019). However, it remains unclear whether these methods will be sufficient for reliably detecting and measuring all types of natural behaviors, such as long-term behaviors involving complex interactions between pairs or groups of individuals, or between individuals and their environments. Alternative strategies may be needed depending on the behavior of interest and the experimental design.

In this study, our primary goal was to automatically measure both developing bower structures and behavioral decisions in naturalistic social environments for extended time periods. We found that a low-cost depth sensor was sufficient for tracking the structural development of bowers over the course of many days, and for capturing natural species differences in bower structure in aquarium tanks that mirror species differences in the wild (York, Patil et al. 2015). Depth sensors have previously been used in behavioral studies, but as tools for animal tracking (Hong, Kennedy et al. 2015, Wiltschko, Johnson et al. 2015). In contrast, we used depth sensing to track the development of an underwater extended phenotype structure in 3D through time. Depth sensors may also be useful for measuring the development of other extended phenotype structures through time such as underwater or above-ground nests, and for tracking activity patterns in animals that construct subterranean structures, e.g. by measuring the volume of substrate that is displaced above ground over time (Theraulaz, Bonabeau et al. 1998, Khuong, Gautrais et al. 2016, DiRienzo and Dornhaus 2017, Genise 2017, Metz, Bedford et al. 2017).

In addition to measuring the bower structure, we also tracked behavioral decision-making on much shorter timescales using action recognition. To our knowledge, this is the first time action recognition has been used to identify and measure complex behaviors in non-human animals. Previous machine learning strategies have classified animal behaviors through analysis of positional tracking and/or pose estimation data (Anderson and Perona 2014, Hong, Kennedy et al. 2015, Robie, Seagraves et al. 2017). In contrast, we rooted our approach in the identification of sand change events from video data, and in doing so we were able to identify tens of thousands of behavioral decisions per trial without tracking or pose estimation. Similar approaches using analysis of background changes at different timescales may be useful for identifying and measuring other construction and/or navigation behaviors defined in part by physical contact and/or interaction with their environments. Similarly, action recognition may be an effective alternative for identifying and measuring a wide variety of natural behaviors in different systems and experimental designs, either in the absence of or in conjunction with positional tracking and/or pose estimation data.

We showed that a 3D ResNet classified video clips of sand change into ten categories with accuracy comparable to a human observer. Remarkably, the model distinguished bower scoops from feeding scoops, and bower spits from feeding spits, despite these behaviors being frequently indistinguishable to an untrained observer. The high prediction accuracy for these behaviors suggests that action recognition may be a powerful tool for studying the evolution of both bower/nest construction behaviors and feeding behaviors in other sand-dwelling cichlid and teleost species. Similarly, high prediction accuracy for quivering, a conserved and stereotyped sexual behavior expressed by many teleosts, suggests that action recognition may be useful for tracking social and mating behaviors broadly across many species, and potentially in other systems in which animals exhibit complex stereotyped behavioral sequences (e.g. courtship behavioral sequences, or aggressive displays). In combination with action recognition, we also applied a Faster-RCNN for object recognition to identify and count fish across behavioral contexts. Notably, both methods achieved high accuracy across three species and one hybrid cross after analyzing relatively small training sets of top down clips/frames, suggesting these are likely adaptable to many other cichlid (and potentially teleost) species and behavioral paradigms utilizing a top-down FOV. Integrating action recognition, object detection, positional tracking, and pose estimation may allow for rich quantitative descriptions of long-term behaviors in many natural systems.

A major strength of our system is the integration of two orthogonal methods to simultaneously measure a developing extended phenotype and the underlying behavioral decisions throughout construction, and the ability to spatially and temporally align these two data streams to quantify relationships between structure and goal-directed decision-making. By analyzing the combined data, we show natural species differences in the relationships between behavior and structure: pit-digging species perform far more scoops in bower regions, and the number of scoops predicts the volume of structural change in these regions; while castle-builders perform far more spits in bower regions, and the number of spits predicts the degree of depth change in these regions. By linking thousands of individual behavioral decisions to a dynamic 3D surface, future studies can dissect the organizing principles through which the developing bower structure modulates decision-making over long timescales.

These methods allowed us to gain new insights into bower construction behaviors that would have been difficult or impossible to achieve through manual analysis. By analyzing depth change through time, we showed that the ultimate bower structure arises through punctuated bursts of activity, typically spanning only a small proportion of daylight hours. This is consistent with field observations in which males leave their bowers for extended periods of time to feed (McKaye 1983). We further show that males construct bowers in a spatially non-uniform manner, exhibiting shifts in spatial patterns of construction over the first three days of building. We also show that males construct bowers in spatially repeatable locations across multiple trials, consistent with observations and studies in the field, in which bower have been experimentally manipulated or destroyed by turbulence from storms, and males reconstruct their bowers with spatial fidelity although not typically in the exact same spatial location (Kirchshofer 1953, Fryer and Iles 1972, McKaye, Louda et al. 1990). Taken together, these data support a role for spatial memory in bower construction, but suggest that a simple, constant, and uniform spatial decision-making program based solely on spatial location is not sufficient for explaining the full trajectory of construction. One possibility is that spatial location drives the male’s decisions about where to initiate construction on a flat sand surface, and as the bower becomes visually salient, physical features of the structure play a more dominant role in modulating decision-making.

We also use depth sensing to demonstrate the sequential expression of pit-digging and castle-building behavior in two pit-castle F_1_ hybrid crosses. The two crosses were made in reciprocal directions (castle-building sire versus pit-digging sire), suggesting that this behavioral sequence is expressed regardless of the sire’s behavioral phenotype. In a previous study, York et al. found a large set of genes exhibiting imbalanced expression of parental alleles in the brain during pit-digging versus castle-building in MCxCV F_1_ hybrids, such that the pit-digging (CV) parent alleles were upregulated during pit-digging, and the castle-building (MC) parent alleles were upregulated during castle-building (York, Patil et al. 2018). Identifying the neuronal populations in which these parental alleles are expressed, and understanding the causal relationships between neural circuits, context-dependent allele-specific expression, and bower construction behavior are important targets for future study.

Integrating action recognition and object recognition also allowed us to gain new insights into behavioral and social dynamics across whole trials. Clear behavioral contexts emerged from temporal analysis of action recognition data, corresponding to feeding, constructing, and spawning contexts. The weak correlations between bower construction and spawning behaviors were surprising to us, given that these are both courtship behaviors. This temporal uncoupling suggests that bower construction and spawning behaviors are triggered by at least partially independent mechanisms, perhaps by differences in visual/chemosensory cues emitted by gravid females and/or differences in the male’s hormonal and neuromodulatory state during spawning. To gain deeper insight into these behavioral contexts, we used object recognition to measure the number of fish present over the sand tray during different categories of behavioral events. We found that social dynamics varied strongly across feeding, construction, and spawning contexts. The number of fish present over the sand tray was lowest during construction behaviors, greater and highly variable during feeding behaviors, and greatest (∼2) during spawning behaviors. This is consistent with spawning occurring in a spatially exclusive manner between the subject male and a single gravid female. The low fish counts during bower construction behaviors are consistent with males aggressively chasing away both male and female conspecifics while constructing and establishing territory prior to spawning, a phenomenon we have previously observed but not quantified in both stock tanks and behavior tanks. An alternative explanation is that females actively avoid the bower during construction. Future analyses of male-female chasing and other aggression behaviors can reconcile these models.

There are several limitations to these experiments that can offer guidance for future development of this paradigm as well as other systems. First, in this study we sacrificed temporal resolution for improved spatial resolution of depth data. Depth sensing with high temporal resolution can be used as a powerful tool for tracking animals against visually complex backgrounds, across 3D trajectories, and/or through occlusions (Anderson and Perona 2014, Dell, Bender et al. 2014, Hong, Kennedy et al. 2015, Wiltschko, Johnson et al. 2015), and thus may be critical for the success of other paradigms. Although sacrificing temporal resolution allowed us to recover a large amount of depth data, the version of the Kinect still yielded a significant degree of data loss. Many new depth sensors with improved time-of-flight technology have been released (including the Kinect v2), but these require USB 3.0 which is not a feature of the Raspberry Pi used in this study (Raspberry Pi 3 Model B+). However, Raspberry Pi has recently released the Raspberry Pi 4, which includes USB 3.0 among other upgrades, opening the door to higher quality depth data and improved temporal resolution at relatively low cost. Another limitation is the practical challenge of remotely controlling a large set of computers and storing, transferring, and analyzing large volumes of video and depth data. For our project, this required planning with information technology professionals at our institution and a Business Dropbox Account for data storage, as well as computer science expertise for developing analysis pipelines. However, these hurdles will likely become less prohibitive as performance specifications improve on low-cost computer systems and more open source and user-friendly computational tools are made publicly available. A final limitation is that our system currently analyzes all video and most depth data after it is collected. Further improvements are needed to enable real-time processing of data, which may be necessary for some projects.

Despite these limitations, these experiments are a significant step for computational ethology, overcoming several major challenges facing the automated measurement of natural long-term behaviors in the lab. Our recording system enables automated phenotyping of naturally evolved construction behaviors in multiple wild-derived species, in naturalistic social environments, over extended time periods. Bower construction behaviors are expressed by more than 200 cichlid species spanning multiple lakes, and an even larger number of species feed in the sand. Our system thus lays a foundation for studying the biological basis of vertebrate behavioral evolution on large comparative scales in the lab. The system is also effective for behaviorally phenotyping interspecies hybrids and will thus be useful for investigating the transition between two species-divergent behaviors in F_1_ hybrids, and for genetic mapping of behavioral variation in F_2_ hybrids. The ability to phenotype many behaviors, and to track thousands of spatial decisions over extended time periods also makes this system particularly promising for future neural recording experiments.

## Conclusions

We have designed, developed, and implemented a behavioral paradigm and recording system for automatically phenotyping construction behaviors in naturalistic social environments in cichlids. By integrating depth sensing and action recognition, we track developing bower structures and decision-making trajectories in multiple species and hybrid crosses over weeklong periods in many tanks simultaneously. This system will help accelerate comparative behavioral genetics and neuroscience experiments in one of the most powerful vertebrate systems for studying natural behavioral evolution.

## Acknowledgements

We would like to thank Tucker Balch for suggestions on tank setup. We thank Andrew Gordus for helpful comments on writing the manuscript. This work was supported in part by NIH R01GM101095 to J.T.S., NIH R01GM114170 to P.T.M., NIH F32GM128346 to Z.V.J., and by Georgia Tech Graduate Research Fellowships to J.L., V.A., and M.A.

## 3. METHODS

### 2.1 Animals and husbandry

#### Subjects

Lake Malawi bower-building species (*Copadichromis virginalis, Tramitichromis intermedius, Mchenga conophoros*) derived from wild-caught stock populations, as well as genetically hybrid individuals derived from these species (described below), were housed in social communities (20-30 individuals) in 190 liter glass aquaria (90.2 cm long x 44.8 cm wide x 41.9 cm tall) into adulthood (>180 days). Aquaria were maintained under conditions reflective of the Lake Malawi environment: pH=8.2, 26.7°C water, and a 12 h:12 h light:dark cycle with 60-minute transitional dim light periods. For all behavioral experiments, a single reproductive adult male and four reproductive adult stimulus females of the same species or hybrid background were introduced into designated home tanks (as described above) equipped with additional LED strip lighting (10 h:14 h light:dark cycle synced with full lights on), and a custom-designed hollow acrylic case (43.1 cm long x 43.1 cm wide x 10.2 cm tall, with a 35.6 cm diameter circular opening) surrounding a circular plastic tray (35.6 cm diameter x 6.4 cm deep, and elevated 3.8 cm above the aquarium bottom) filled with sand (Carib Sea; ACS00222). Sand trays were positioned approximately 58 cm directly below a Microsoft XBox Kinect depth sensor and Raspberry Pi video camera; and approximately 30 cm directly below a custom-designed transparent acrylic tank cover (38.1 cm long x 38.1 cm wide x 4.4 cm tall) that contacted the water surface to eliminate rippling for top-down depth sensing and video recordings (described below). In both stock and behavioral tanks, fish were fed twice daily with dried spirulina flakes (Pentair Aquatic Eco-Systems).

#### In vitro hybridization

Reproductively active males and females were visually identified based on abdominal distension (females), nuptial coloration (males), and expression of classic courtship behaviors (e.g. chasing/leading and quivering). Two separate pit-castle hybrid crosses were generated in the reciprocal direction: *Tramitichromis intermedius* (female) x *Mchenga conophoros* (male); and *Mchenga conophoros* (female) x *Copadichromis virginalis* (male). To cross-fertilize, a petri dish was filled with water from the home tank, and eggs were collected into the dish by applying gentle pressure between the pectoral region and the anal pore of the female. Eggs remained fully submerged while the male’s sperm was extracted into the same dish by applying gentle pressure to both sides of the abdomen. The mixture was immediately and gently agitated and then eggs were gently rinsed twice with fresh aquarium water to reduce polyspermy. Eggs were then transferred into a beaker containing a fresh oxygen tube, fresh aquarium water, and a drop of methylene blue to minimize risk of fungal infection. Water replacement was performed at least once daily until hatching (approximately 5-6 days post-fertilization).

#### Behavioral trials

For each behavioral trial, a single reproductive adult subject male was introduced to a designated behavioral tank containing four reproductive adult stimulus females and a full sand tray as described above (under Animals and husbandry). Upon introduction, an automated recording protocol (described in detail below) was initiated, collecting RGB video and depth data during full light hours (08:00 to 18:00 EST) for 7-10 days. Subjects and stimulus females were allowed to freely interact throughout the entirety of the recording trial and followed the same feeding schedule described above (under Animals and husbandry).

### 2.2 Recording and monitoring system

#### Hardware

The automated recording system consisted of a Raspberry Pi 3 Model B (RASPBERRYPI3-MODB-1GB; Raspberry Pi Foundation) connected to the following: (1) a 7” touchscreen display (RASPBERRYPI-DISPLAY; Raspberry Pi Foundation) secured in an adjustable mount case (Smarticase); (2) an Xbox 360^®^ Kinect™ Sensor (Microsoft); (3) a Raspberry Pi camera v2 (RPI 8MP CAMERA BOARD; Raspberry Pi Foundation); and (4) a 1 TB external hard drive (WDBUZG0010BBK-WESN; Western Digital).

#### Code

We wrote custom Python scripts for all aspects of the project. All code is publicly available on github at www.github.com/ptmcgrat/Kinect2. A general outline of the code is available in the Supplementary Materials.

#### Depth sensing

We used a Microsoft Xbox Kinect depth sensor to measure the topology of the sand surface through water. The Kinect is a low-cost, close-range, high-resolution depth sensor containing an IR laser and refractor that emits a known structured light pattern, and an IR camera that detects the emitted IR light across surfaces within the FOV. The Kinect then uses a pattern recognition algorithm to compute distance of surfaces across the FOV, which can be stored into 640×480 numpy array files (.npy). Kinect depth sensing was controlled through a custom Python script (the CollectData function within the CichlidBowerTracker.py script, see Supplementary Materials and Methods) that was initiated at the beginning of each behavioral trial. Because continuous depth data was both unnecessary and impractical (due to the large volume of high frame rate uncompressed depth data), CollectData combined depth data collected continuously at ∼10 Hz into a single frame every 5 minutes. The code also specifies collection of a single RGB snapshot every 5 minutes, for later registration between depth data and video data. All depth data was stored on an external hard drive for later processing.

#### Video recording

The same CichlidBowerTracker.py script controlled daily collection of 10 hours of 1296×972 RGB video through a Raspberry Pi v2 camera (Raspberry Pi Foundation), data during full lights on hours (08:00-18:00 EST). The large volume of video data collected per day was enabled by instantaneous compression into .h264 format by the Raspberry Pi. Compressed video data was stored on an external hard drive for later processing.

#### Google Controller spreadsheet

A Google Controller spreadsheet was created to remotely control each tank’s Raspberry Pi recording system, provide real-time visual updates of bower activity every five minutes, and logging behavioral trial information into a master datasheet. The Controller sheet served as a master graphical user interface for the recording system, with a “Command” column monitored by each Raspberry Pi. The Commands included “New” to initiate a new trial, “Restart” to resume an existing trial, “Rewrite” to overwrite an existing trial, “Stop” to stop a trial, “UploadData” to upload data from a completed trial to Dropbox, and “LocalDelete” to clear data from the local storage drive following upload. A more detailed description of Google Controller setup and functionality is provided in the Supplement (subsection “Controller Spreadsheet”).

### 2.3 Data processing and analysis pipeline

#### Data upload

Following completion of each trial, data was copied from the local external hard drive to a laboratory Dropbox account through the Google Controller spreadsheet by upload through rclone, a cloud storage sync program (https://rclone.org/). The directory for each trial contained all videos, RGB frames, and depth frames recorded for the trial. Due to the large volume of data, uploading for all data collected in a recording round (∼10 trials, ∼4TB of data) typically required 24-28 hours. For later trials, upload time was reduced to ∼3-5 hours by first compressing depth data into .tar files.

#### Depth Analysis

Analysis of Kinect depth data for each trial was performed using the DepthAnalysis function within the DataAnalyzer module of the CichlidBowerTracker.py script. Depth analysis included the following: (i) conversion of raw depth data to “millimeters from Kinect”, (ii) smoothing depth data by applying a Savitsky-Golay filter to spatial and temporal dimensions of raw depth data using the savgol function in Python, (iii) frame-to-frame subtraction (and visualization) of smoothed data at whole trial, daily, and hourly timescales, (iv) identification of above-threshold depth change (whole trial: ±1.0 cm, daily: ±0.5 cm, hourly: ±0.18 cm) regions at each of these timescales, (v) identification of the single highest change region (bower ROI) at each of these timescales, and (vi) calculation of several indices of structural change at these timescales: pixel size of above-threshold depressed (pit-like) and elevated (castle-like) regions; volume of above-threshold depressed (pit-like) and elevated (castle-like) regions; and four calculations of the “Bower Index” (the net volume change divided by the absolute volume change): the overall Bower Index for all depth change, and three for above-threshold change only using sequentially increasing depth thresholds (Trial: 1.0 cm, 3.0 cm, 5.0 cm; Day: 0.4 cm, 0.8 cm, 1.2 cm; 2-hour: 0.2, 0.4, 0.8). The final Bower Index used for analyses was the average of these four calculations.

#### Video Analysis

Analysis of sand change in video data for each trial was performed using a custom VideoProcessor.py script. VideoProcessor.py includes the following: (i) a Hidden Markov Model (HMM) algorithm to detect changes in pixel values through time, and (ii) a density-based clustering algorithm to identify clusters of HMM+ pixels, or putative sand change events. Briefly, for each video, the value of each pixel was analyzed through time, and a custom HMM algorithm was used to predict enduring changes in pixel values using the ‘hmmlearn’ package for Python. This script simultaneously ignored short-term changes that could be caused by fish swimming. To improve computational efficiency, pixel values were sampled at a rate of 1 value per 30 frames (equivalent to once per second). This analysis generated a 3D sparse matrix in which “0” represented no change and “1” represented HMM-predicted change. Because some of the HMM-predicted changes could be caused by noise (e.g. variance in pixel value caused by the camera sensor) we used density-based spatial clustering of applications with noise (DBSCAN) within the Python package ‘sci-kit learn’ to identify clusters of HMM+ change in the presence of noise. DBSCAN parameters were set based on observed size of sand change events and from a k-dist graph (see Supplementary Methods subsection “Density-based clustering for identification of putative sand change events”). DBSCAN analyzed each HMM+ pixel change point in time and space, and used a KD-tree to determine if the neighboring region contained a minimum number of HMM+ points. This enabled us to identify spatiotemporal clusters of HMM+ pixels, representing putative sand change events.

### 2.4 Machine Learning

#### Deep learning of cichlid behaviors

For each cluster of putative sand change pixels identified by density-based clustering, a four second (120 frame) 200×200 pixel RGB video clip was generated, centered spatially and temporally around the sand change event. A trained observer manually classified 14,234 video clips randomly selected from representative days across seven trials, spanning seven subjects, three species, and one hybrid cross. Each clip was classified into one of the following ten categories: bower scoop, bower spit, bower multiple event, feeding scoop, feeding spit, feeding multiple event, spawning, drop sand, other-fish, and other-no fish. The operating definition for each behavioral category is provided in the Supplementary Material (subsection “Behavioral definitions for manual annotation”). We used 80% of manually annotated clips for training an18-layer 3D ResNet, and the remaining 20% of clips were used for testing. Briefly, 3D ResNets are 3D convolutional neural networks (CNNs) that incorporate features of Residual Networks (ResNets), in which signals are bypassed across convolutional layers during training. This approach allows 3D ResNets to be deeper and more accurate than traditional 3D CNNs for action classification tasks (Qiu, Yao et al. 2017). For training, testing, and prediction we used the 18-layer architecture described in (Qiu, Yao et al. 2017) (https://github.com/kenshohara/3D-ResNets-PyTorch). Prior to training and testing, each video clip was first converted to 120 RGB images in .jpeg format using ffmpeg, and during training images were randomly cropped at multiple scales and resized to 112×112 pixels per image, and then randomly flipped at a rate of 0.5 for data augmentation. Each channel was then normalized based on the mean value for that channel across all videos. For training, stochastic gradient descent was used to optimize the parameters of the neural network. Specifically, the learning rate was set to 0.1 (and set to decrease after 10 consecutive epochs of no change in loss), momentum was set to 0.9, dampening was set to 0.9, weight decay was set to 1.0×10^−4^. The network was trained for 100 epochs with a batch size of 8 per epoch.

#### Deep learning for fish detection and counting

To detect and count fish we used a Faster region-based convolutional neural network (Faster-RCNN). Faster-RCNNs are two-step neural networks for fast and accurate object detection. In the first step, a pretrained convolutional neural network (CNN; in these experiments a ResNet50 trained on the COCO dataset, http://cocodataset.org). extracts features from the raw image, and then these features are fed into a Region Proposal Network (RPN) which identifies ROIs that may contain objects of our interest. In the second step, these ROIs are analyzed by a convolutional neural network which classifies objects of interest and generates bounding boxes using linear regression. Our dataset consisted of 1842 manually annotated frames sampled from seven trials. Fish in each frame were annotated using labelImg (https://github.com/tzutalin/labelImg) and the annotations were stored as .xml files. We used Tensorflow models (https://github.com/tensorflow/models) to preprocess data. 80% of the dataset was used as a training set (n= 1473) and the remaining images were used as the test set (n= 369). Manual annotations were then used to train both RPN, CNN classifier and bounding box regressor.

### 2.5 Statistics

All statistics were performed using Python 3 (version 3.6 or later) and R (version 3.4.4 or later).

#### Depth change by condition

Sand displacement between conditions was compared using the with whole trial depth change as the outcome variable and condition (empty tank trials vs. “no bower” control trials vs. bower trials) as the predictor variable. We tested the assumption of heterogeneity of variance using the Fligner-Killeen test, which revealed unequal variance among groups. Based on this, we tested differences between groups using the Kruskal-Wallis H test (non-parametric one-way ANOVA on ranks). Post-hoc pairwise Wilcoxon Rank Sum Tests were performed to assess pairwise significance among groups.

#### Depth change thresholds

To identify depth change thresholds, we quantified whole trial, daily, and hourly depth change across a large and representative sample of control (n=22) and bower (n=27) trials. To filter out signals due to noise, we set a minimum size threshold of 1,000 contiguous pixels (∼10 cm^3^) using remove_small_objects within the morphology module in the scikit-image library for Python (for example usage see DepthProcessor.py code at https://github.com/ptmcgrat/Kinect2/blob/master/Modules/Analysis/DepthProcessor.py). We then incrementally applied depth change thresholds in 0.1 mm steps to identify the maximum values that could be expected in the absence of bower construction. These thresholds turned out to be 1.8 mm for hourly change, 5.0 mm for daily change, and 10.0 mm for whole trial change.

#### Bower Index by species

The Bower Index was calculated as the sum of above threshold depth change (directional; positive and negative changes cancel out) divided by the sum of total depth change (absolute value; change in either direction is considered positive) at each timescale. To account for variation in building intensity between individuals, we applied stepped increases in the depth threshold at each timescale, and we averaged together the bower indices calculated using each threshold. Bower indices were compared between species (MC vs. CV vs. TI) using one-way ANOVA and significance of pairwise comparisons were analyzed with post-hoc Tukey’s HSD tests.

#### Spatial repeatability

To measure spatial repeatability we analyzed all above threshold pixels in each paired trial. The percentage of spatial overlap was calculated as the proportion of these pixels that was above threshold in the same direction in both trials. To determine whether spatial overlap was greater than overlap expected by chance, we calculated the expected overlap for depressed regions and elevated regions independently (pits can be dug within the sand tray region, but not in the acrylic platform, whereas castles can be built within the sand tray region or on the acrylic platform). To determine the overlap expected by chance for depressed regions, we calculated the proportion of the sand tray that these regions occupied in each paired trial, and multiplied those proportions together. To calculate the overlap expected by chance for elevated regions, we calculated the proportion of the sand tray and platform that these regions occupied in each paired trial, and multiplied those proportions together. The total expected spatial overlap was calculated as the sum of these two numbers. We used paired t-tests to analyze whether the degree of spatial overlap observed in repeatability trials and control paired trials was greater than expected by chance. To test whether spatial repeatability was greater within subjects than between subjects, we analyzed males that were tested in the same tank as other males of the same species (n=7 total; CV, n=3; MC, n=1; TIxMCF1, n=2; MCxCVF1, n=1). For each subject, we took the average spatial overlap with other males of the same species, and compared it to the actual spatial overlap observed between repeatability trials using a paired t-test.

#### Spatial uniformity

To calculate the Spatial Uniformity Index, we first measured the whole trial volume change as the sum of daily above-threshold volume changes. We then defined the whole trial region of interest as the union of all daily above-threshold regions. We defined the final structure as the whole trial depth change within the whole trial region of interest. To estimate expected volume change, we first calculated an expected change ratio as the ratio of daily above-threshold volume change to whole trial volume change. To calculate the expected volume change on each above-threshold day, we multipled the final structure by the expected change ratio for that day. By taking the difference between the expected depth change map and the actual depth change map, we were able to quantify how structural developments diverged from spatial uniformity with a Spatial Uniformity Index (SUI):

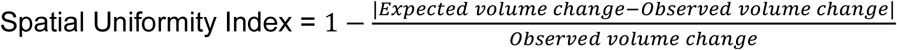

For primary analyses of spatial uniformity we used a linear mixed-effects regression model with SUI as the outcome variable; species, day, and the interaction between species and day as fixed effects; and subject as a random effect. Thus the model was as follows:

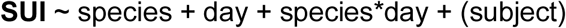

We performed an additional analysis to test whether accounting for daily volume change would significantly alter our results. Thus, we included daily volume change as an additional fixed effect in the model:

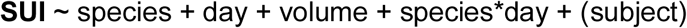

For all linear mixed effects models, we calculated estimates for fixed effects by maximum likelihood estimation using the ‘lme4’ package in R and calculated significance for fixed effects using Satterthwaite approximation through the ‘lmerTest’ package and the ‘anova’ function in R. Estimates of pairwise differences between levels for each fixed effect were calculated using estimated marginal means (least squared means), and the significance of these differences were determined using Satterthwaite approximation corrected for multiple comparison families with Tukey’s adjustment, using the ‘emmeans’ and ‘multcomp’ packages in R.

#### Behavioral correlation analysis

To quantify patterns of temporal covariance among different behavioral categories and total depth change, we performed correlation analyses on all seven trials that were analyzed by the 3D ResNet. Each trial was divided into 60-minute time bins. Within each 60-minute bin, the number of events was calculated for each category, as well as the total absolute volume change from depth data, and times bins in which no behavior from any category were expressed were excluded from analysis. Pairwise behavior-behavior and behavior-depth change correlations were then performed across all bins and trials (pooled).

#### Fish counts across behavioral contexts

To count fish across different behavioral contexts we extracted whole frames from four second time periods associated with 3D ResNet-predicted behavioral events. We predicted fish locations as well as the total fish count in each frame. We then calculated the average fish count for each event. To analyze differences in number of fish present in frames associated with different behavioral contexts, we used a linear mixed-effects model with fish count as the outcome variable, behavior as a fixed effect, and day nested within subject nested within species as a random effect. Thus the model was as follows:

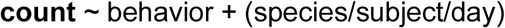

Estimates of counts by behavior, differences in count between behaviors, and the significance of these differences were calculated using the same methods as described above under “Spatial Uniformity”.

#### F_1_ behavior through time

To analyze the Bower Index in pit-castle F_1_ hybrids through time, we used a linear mixed-effects model with Bower Index as the outcome variable; cross, day, and the interaction between cross and day as fixed effects; and subject as a random effect:

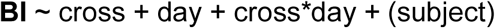

Associations between Bower Index and day, cross, and the day*cross interaction were calculated as described above. Differences in count between behaviors, and the significance of these differences were calculated using the same methods as described above under “Spatial Uniformity”. Post-hoc comparison of the Bower Index across days in F_1_ hybrids with the Bower Index across days in hybrids was performed using one-way tests.

## Supplementary Methods and Materials

### System Design

Animal care guidelines required that testing over such extended time periods had to be done in the home tank (as opposed to external testing arenas). In our facilities, home tanks are supported on tank racks with built-in piping and support beams that partially occlude top-down fields of view (FOVs) (e.g. see Supplementary Figure 1). Additionally, all tanks have a central support crossbeam that partially occludes top-down FOVs. We found that a ∼36 cm diameter sand tray placed on one half of the home tank provided a sufficient volume of sand for males to construct bowers, and was small enough to fit into an unobstructed top-down FOV (Supplementary Figure 3B). We designed a custom acrylic platform to surround the sand tray to prevent subjects from spitting sand over the edge of the tray onto the bottom of the aquarium. Thus, in this design subject males and females could freely enter and exit the sand tray region throughout the trial.

**Supplementary Figure 1.**
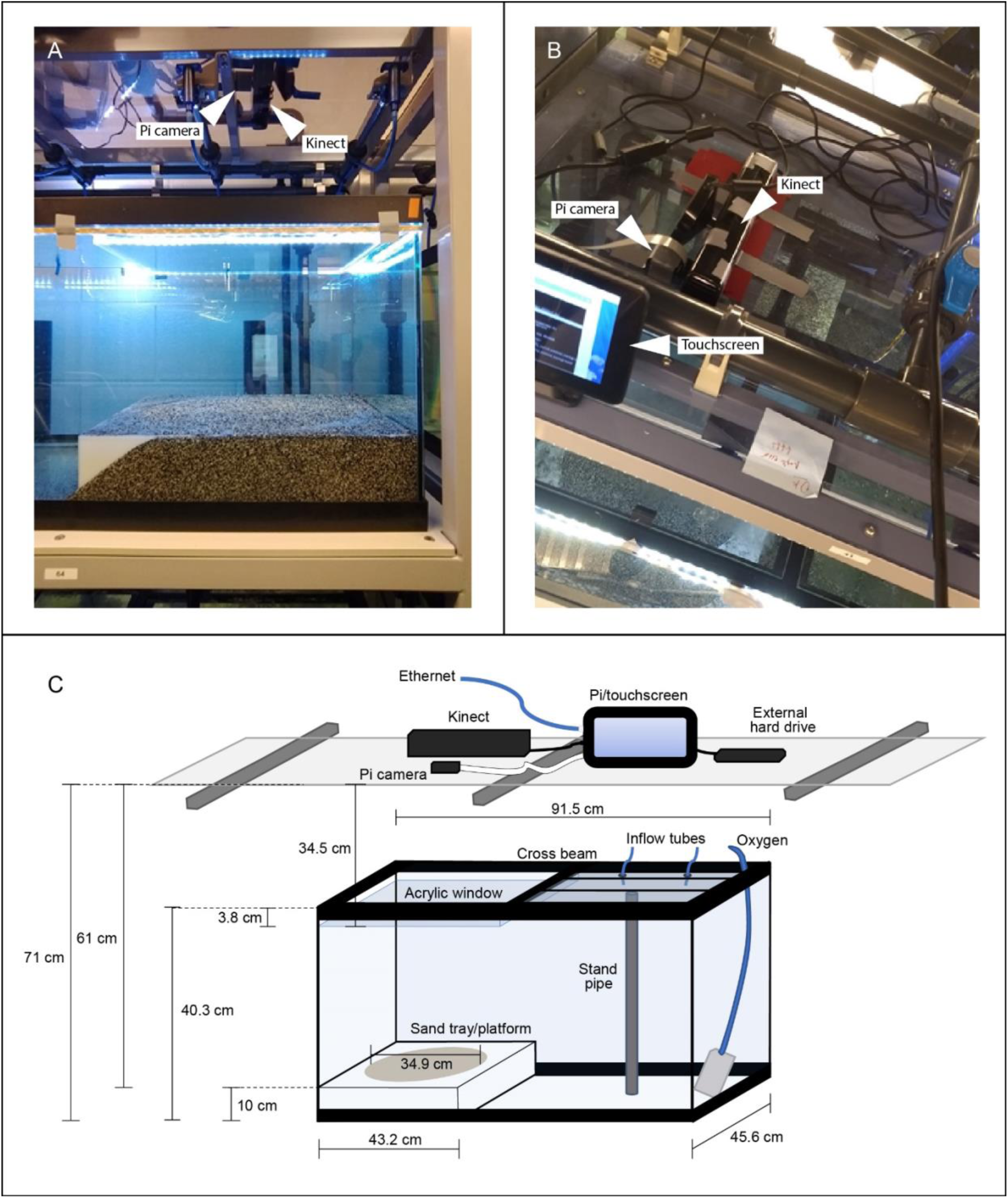
Photographs, schematic, and measurements of behavioral paradigm. Photographs (A-B) and detailed dimensions of home tank setup for bower behavior assays (C). The final design had to be compatible with several pre-existing physical constraints such as tank rack support beams (gray metal beams visible just beneath acrylic in A, B), water inflow lines (gray acrylic and blue rubber tubes above and below transparent acrylic top, visible in A and B), and aquarium cross beams (black plastic cross beam visible in B). All electronic equipment was placed on top of a transparent acrylic shelf above the tank rack, with the Kinect and Raspberry Pi camera (indicated with white arrows in A, B) aimed downwards for a top-down view of the sand tray.

**Supplementary Figure 2.**
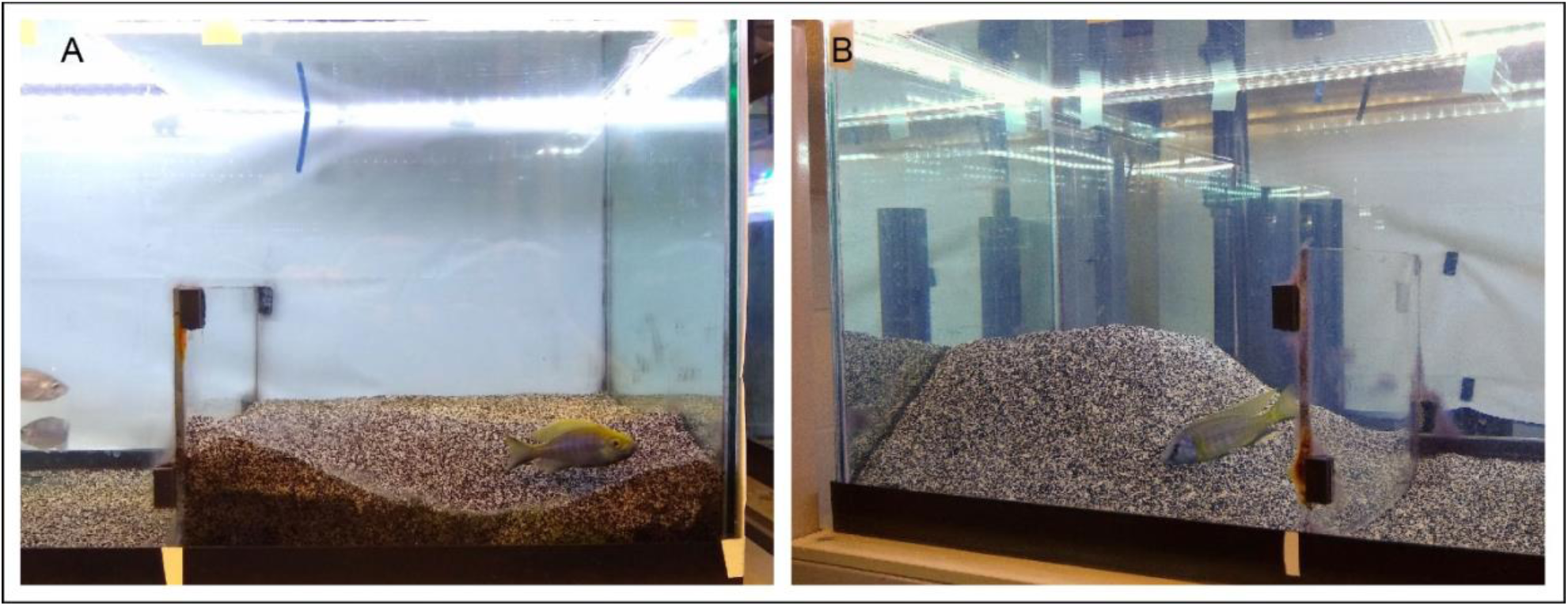
Photographs of bower structures in the lab. Photographs of a *Copadichromis virginalis* male and his pit (A) and of a *Mchenga conophoros* male and his castle (B) in a modified behavioral tank setup.

### Controller Spreadsheet

To avoid the need for manual control of recording equipment above behavior tanks, we created custom software to remotely control each unit using a single Google Spreadsheet: each Raspberry Pi monitored one of the rows of the spreadsheet for commands (Record, Rewrite, Stop, etc.) and executed accordingly (Supplementary Figure 3). The Pi also continuously forwarded analyses of depth change over the previous hour, day, and whole trial to the Google spreadsheet for remote visualization of bower activity (Supplementary Figure 4). This system thus allows for real time monitoring of bower construction.

To setup the Google spreadsheet, two different Python APIs were used to easily access Google APIs: Gspread and PyDrive. Gspread is a module that specifically manages Google Spreadsheets, while Pydrive manages files more generally in Google Drive. In our setup, PyDrive was used to upload .jpeg files containing snapshots and summary images to Google Drive, and Gspread was used to read and write directly to the Controller sheet. The latest documentation and downloads for Gspread are available here: (https://gspread.readthedocs.io/en/latest/index.html) and for Pydrive here: (https://pythonhosted.org/PyDrive/#).

A new Google account was created to house the Google Spreadsheet. We recommend for several reasons. First, this limits the possible exposure of a personal Gmail account since different authentication keys or tokens will need to be distributed to each system that requires access. Second, a new account may also be useful if an automated email system is implemented because it can act as the originating email address that all Pi systems can access.

All authentication for Google APIs goes through OAuth2, but these two modules require different credentials. Gspread requires a Service Account Key, and Pydrive requires a client secret .json file. The latest instructions on how to obtain these credentials and how to use them for authentication can be found in these modules’ documentations, for (Gspread: https://gspread.readthedocs.io/en/latest/oauth2.html, and for Pydrive: https://gsuitedevs.github.io/PyDrive/docs/build/html/quickstart.html#authentication.

After obtaining the appropriate credentials, each Pi needs to have both Gspread and Pydrive downloaded and installed, the service account key for Gspread, the client secret .json file for Pydrive, and an internet connection. This basic setup can easily be customized to fit other experiments in several ways that include but not are limited to adding or changing the modules used and changing the organization and information relayed to the Controller Spreadsheet.

### Automated Email System

An automated email system was setup to send summary updates of the current status for all Pi systems at the beginning and end of each day, as well as real-time notifications of when recordings were unexpectedly interrupted. The basic procedure of this python script is to first check the Controller sheet for nonresponsive Pi systems or to check on the status of all the Pi systems for a summary update. The information from this check is stored and then written into an email which is sent through the Google account’s Gmail. To run this procedure, the Python script was run continuously on a single Pi system with internet connection, the Service Account Key for Gspread, and a .txt file containing the username, password, and email addresses of recipients. The essential modules for the script were Gspread for reading into the Controller sheet and smtplib for sending the email. More information about smtplib and an example of how to use this module can be found here: https://docs.python.org/3/library/smtplib.html.

**Supplementary Figure 3.**
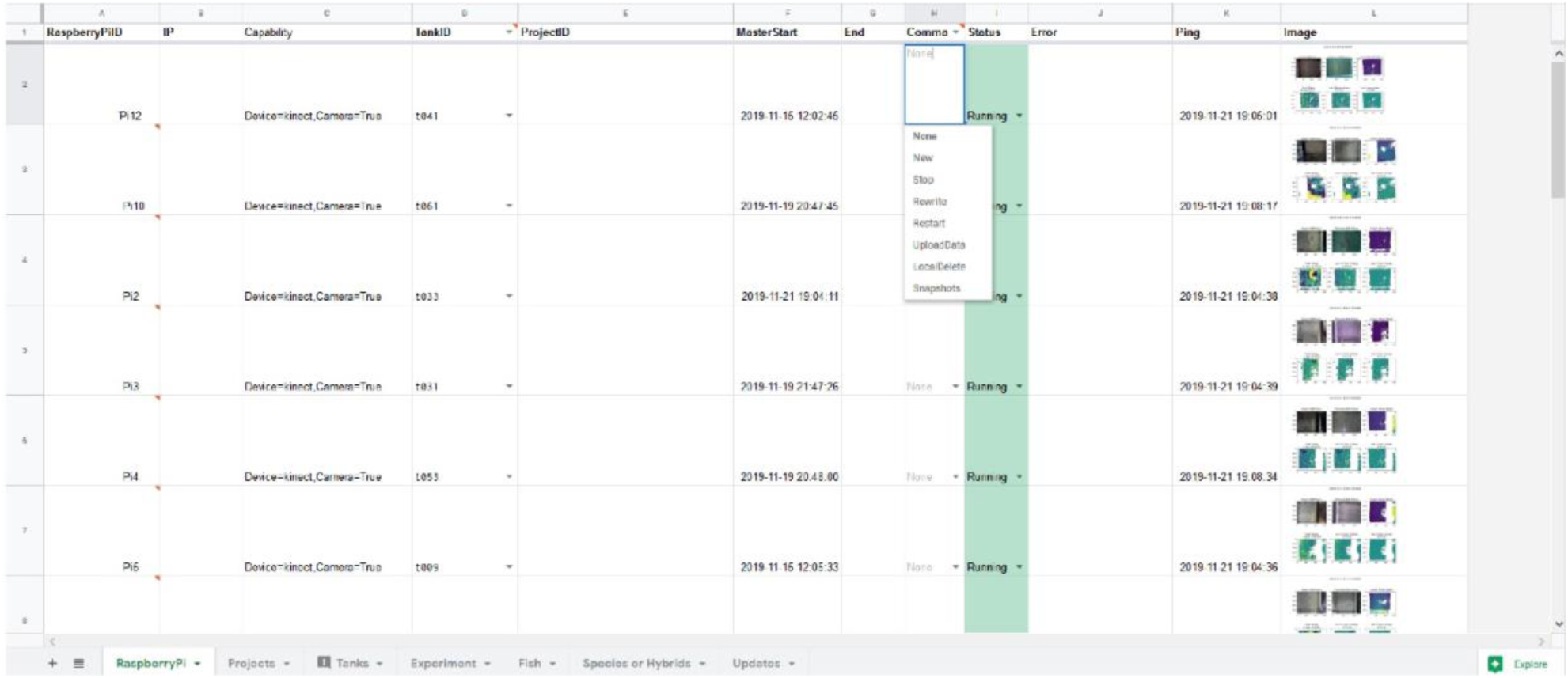
Google Drive Controller spreadsheet for remote control of Raspberry Pi systems. All Raspberry Pi systems were remotely controlled through a Google Drive Spreadsheet. The master spreadsheet comprised multiple sub-sheets for organizing trial information. The first sheet, “RaspberryPi” shown above, was used to remotely issue commands to each Pi unit through a Command Column including Start, Stop, Restart, Rewrite, Upload (to Dropbox), Delete, and Snapshots (shown in blue outlined box above). The current status of each Pi was continuously updated in a separate “Status” column (all green cells reading “Running” indicate actively recording trials). An “Error” column displayed errors encountered during interruptions to help with troubleshooting and debugging. The “Ping” column registered pings from each Pi released every five minutes, and could also be used to identify interruptions. The final “Image” column updates every five minutes provides RGB and depth snapshots to enable live monitoring of depth change across the whole trial, the previous day, and the previous hour.

**Supplementary Figure 4.**
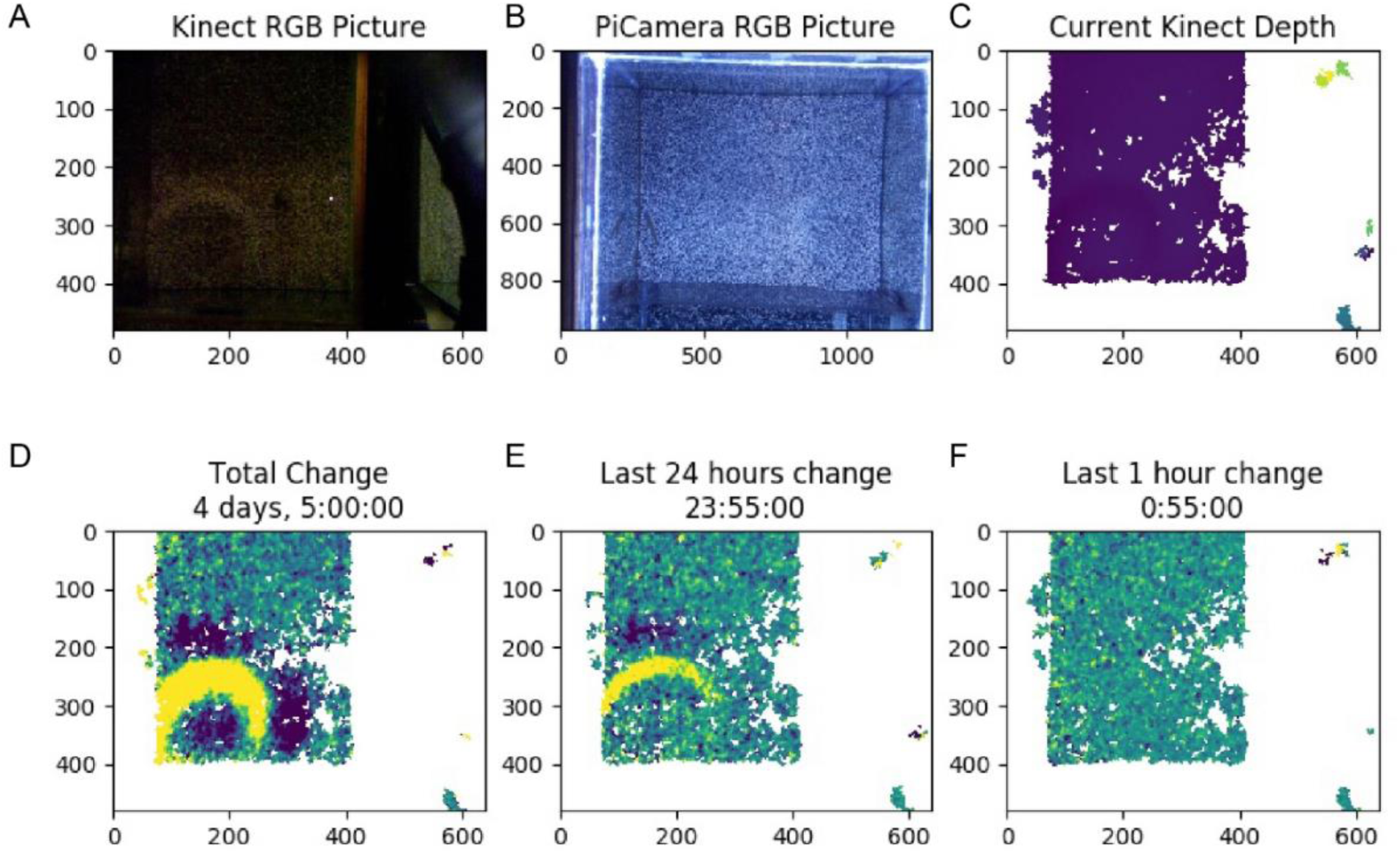
Example screenshot of live update of depth change in behavior tank. Full view of .jpeg file generated every five minutes in the Image column of the Google Controller spreadsheet. The file contains an RGB image captured by the Kinect (A), and RGB image captured by the Raspberry Pi camera (B), the current depth across the sand surface (C), the total depth change across the whole trial as well as the current duration of the trial (D), depth change in the previous 24 hours (E), and depth change in the previous hour (F). Labels on x- and y-axes indicate pixel dimensions.

#### Depth Sensing System Validation

##### The Kinect measures the distance of the sand tray surface through water

The Kinect depth sensor records both depth data and RGB data across the FOV. This sensor was designed for detecting depth changes through air (i.e. Microsoft Xbox users playing video games in their living rooms). In preliminary experiments, we tested how the ∼27 cm of water between the Kinect and the sand tray would interfere with its ability to measure distances of surfaces along the bottom of aquarium tanks. We found that individual snapshots of the sand surface contained a large amount of missing data, potentially due to reflection at the water surface boundary and absorption by water. For example, in a sample set of raw snapshot frames, we found that 40.0 ± 0.04% of pixels per frame contained missing data (Supplementary Fig. 2A). To improve our measurements of the sand surface, we modified our protocol to collect five minutes worth of snapshots in rapid succession (∼10 fps) and average them into a single frame. Although this reduced the temporal resolution of depth sensing, this limitation was reasonable because we expected structural changes of interest to occur over the course of hours. Averaging drastically reduced the number of NaN pixels in each frame (Supplementary Fig. 2B; proportion of NaN values decreased to 20.6 ± 0.06%). We also applied spatial interpolation (see Methods) to estimate values in small regions of missing data, which further reduced the proportion of NaN pixels to 10.8% for the final analyzed dataset (Supplementary Fig. 2C). Thus, our pipeline generated depth data across ∼90% of the sand tray surface every five minutes, enabling analysis of surface change through time.

##### Thresholds improve signal-to-noise for measuring bower construction

We detected significant depth change signals in empty tanks and in control trials, presumably due to noise and other behaviors that alter the sand surface, respectively. Based on these results, we tested if thresholds could separate signals caused by bower construction from signals caused by noise and other non-bower behaviors. We measured the maximum whole trial volume change signals observed in empty tank and control trials (this turned out to be 1.0 cm), and then tested whether volume change signals in bower trials exceeded this threshold. Indeed, we identified greater depth change signals in every bower trial (29/29; Supplementary Fig. 2F), suggesting that threshold could be used to filter out low magnitude depth change signals caused by noise and other non-bower behaviors (Supplementary Fig. 2G).

##### Measurement of bower activity on shorter timescales

We next tested whether bouts of bower activity within trials could be detected on shorter timescales by analyzing depth change over 24-hour and 2-hour periods. We used a similar approach to identify thresholds that separated depth change during bower trials from depth change during control trials. Again, we found thresholds that separated daily and hourly depth change in bower trials versus control trials. Overall, 160/264 (60.6%) of all days analyzed, and 538/3,168 (17.0%) of all 2-hour bins analyzed contained depth change exceeding these thresholds (Supplementary Fig. 2H). Frame-to-frame subtraction of depth data in 5-minute intervals further revealed sharp peaks in activity punctuated throughout whole trials (e.g. see Supplementary Fig. 2I, representative castle-building MC trial).

**Supplementary Figure 5.**
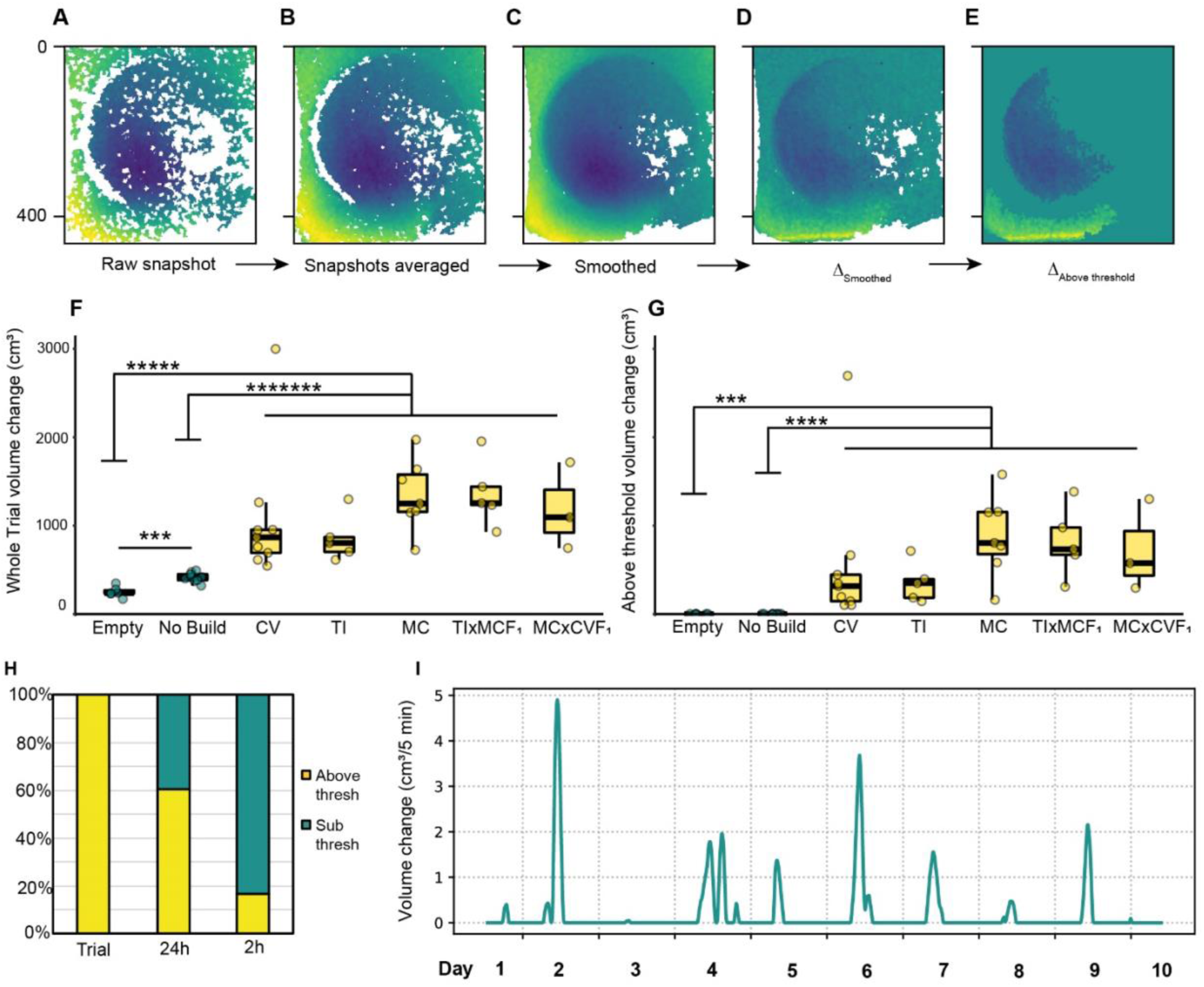
Depth sensing detects height change across the sand surface at different timescales. The Kinect collects top-down depth snapshots of the sand tray surface through time (A-E), with yellow indicating distances closer to Kinect (elevated regions), and dark blue indicating regions farther from the Kinect (depressed regions). Raw Kinect depth snapshots of the sand tray surface contained ∼40% missing data (white pixels; A). To improve depth data quality, consecutive depth snapshots were collected and averaged together every five minutes, reducing the proportion of missing data to ∼20% (B). Data quality was further improved by spatially interpolating data in small NaN “islands,” reducing the proportion of NaN pixels to ∼10% (C). Depth change over the course of the trial was calculated by subtracting the initial depth map from the final depth map, with turquoise indicating no change (D). Thresholding enabled depth change signals caused by bower construction to be separated from signals caused by noise and other home tank activities (E). Before thresholding, total volume change differed strongly between control conditions (Empty tank trials, and trials in which no bower was constructed) compared to bower trials (F). Following thresholding, all bower trials exhibited above threshold volume change while control trials did not (G,H). At shorter timescales, 60.6% of all 24-hour bins analyzed and 16.7% of all 2-hour bins analyzed contained above threshold depth change (H). Analysis of hourly depth change over the course of whole trials revealed that structural change was driven by short bursts of activity punctuated throughout trials (representative *Mchenga conophoros* trial, I).

### Code

All code for the recording system is available at https://github.com/ptmcgrat/kinect2

### Density-based clustering for identification of putative sand change events

A hidden Markov model was used to identify all instances of long-term change in pixel value in each video, and this information was stored as a sparse matrix. From these raw HMM output numpy arrays, we extracted all HMM state changes in a format of (timepoint, y_coordinate, x_coordinate, and state_difference). To separate noise from potential sand manipulation events, we use density-based spatial clustering of applications with noise (DBSCAN) in the Python package sci-kit learn (Pedregosa, Varoquaux et al. 2011). DBSCAN analyzes the region surrounding each HMM+ pixel in time and space, determines if the neighboring region contains a minimal number of HMM+ pixels, expands on dense groups of points, and repeats. DBSCAN parameters were based on estimation from a k-dist graph and observed pixel size of sand change caused by spit and scoop events. A KD-tree was used to quickly and sparsely calculate pairwise distances between sand change points. The clusters were annotated by three human observers independently to assess the quality of events and build a training set for event classification.

#### Pre-processing

The pre-processing workflow output includes cluster identity and coordinates in numpy format, video clips centered spatially and temporally around each cluster for annotation, and histograms and scatter plots to visualize clusters in the video. The workflow also provides options to plot and visualize HMM data before clustering to help set parameters for pre-processing the data. For example, we pre-processed HMM sand change data with the following methods (options included in the script):

1. We filtered out the n timepoints that contained the most HMM+ pixels in a second: to address false-positive sand change signals caused by changes in indoor lighting, this parameter allows a threshold to be set to exclude clusters associated with above-threshold amounts of total HMM+ pixels
2. Thresholding on the magnitude of HMM state difference: some low magnitude changes result from natural variance in pixel values produced by the camera. A threshold for this magnitude can be identified using a histogram of all magnitude of all HMM+ pixel change magnitudes (e.g. see distribution of pixel change magnitudes in Supplementary Figure 4).
3. Masking the tank region: the video can include outer tank walls, reflections, and regions outside the tank entirely. A mask outlining the tank region can be manually drawn to exclude data in regions that are not of interest.

**Supplementary Figure 6.**
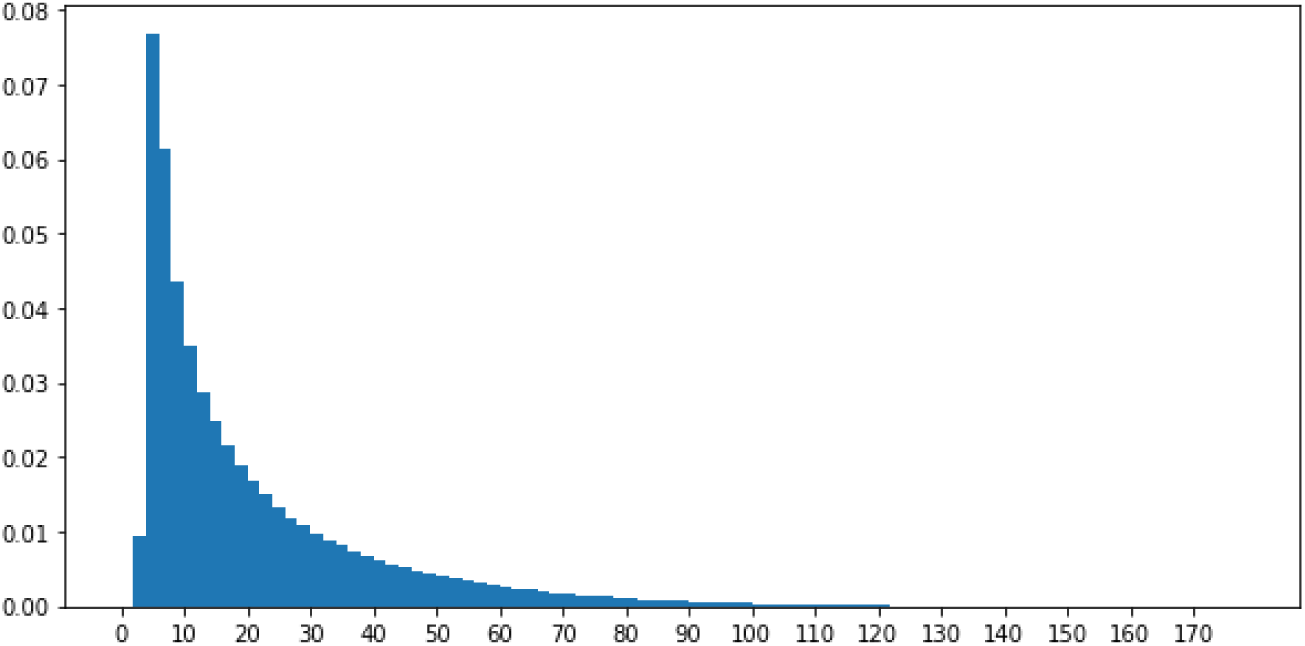
Proportion of HMM+ pixels exhibiting different magnitudes of pixel value change. Distribution of pixel value change magnitudes over the course of a full day of video recording in a representative *Mchenga conophoros* trial.

#### Parameters for density-based clustering

*DBSCAN minPts and eps:*

1. minPts: observers reviewed several hundred putative sand change events and estimated the minimum size of a true sand change cluster to be 10 pixels x 10 pixels x 3 frames, and HMM+ pixels change to cover at least 15% of the putative sand change region. Based on these estimates we calculated the range for the minimum number of pixels in a sand change event to be between 50-250 pixels.
2. eps: For a given k we defined a function k-dist from the database D into the non-negative real numbers, mapping each point to the distance from its k-th nearest neighbor. After sorting the points in the database in descending order based on their k-dist values, the graph of this function suggested a density distribution in the database. This graph is called the sorted k-dist graph, as described in (Ester, Kriegel et al. 1996). We then fit a nearest neighbor tree to all points and use the keighbors query to find the minPts^th^ nearest neighbor for each point, and the k-dist graphs for minPts = 200. We found that most of the points were close to each other; and most points had at least 200 points within 40 units.

**Supplementary Figure 7.**
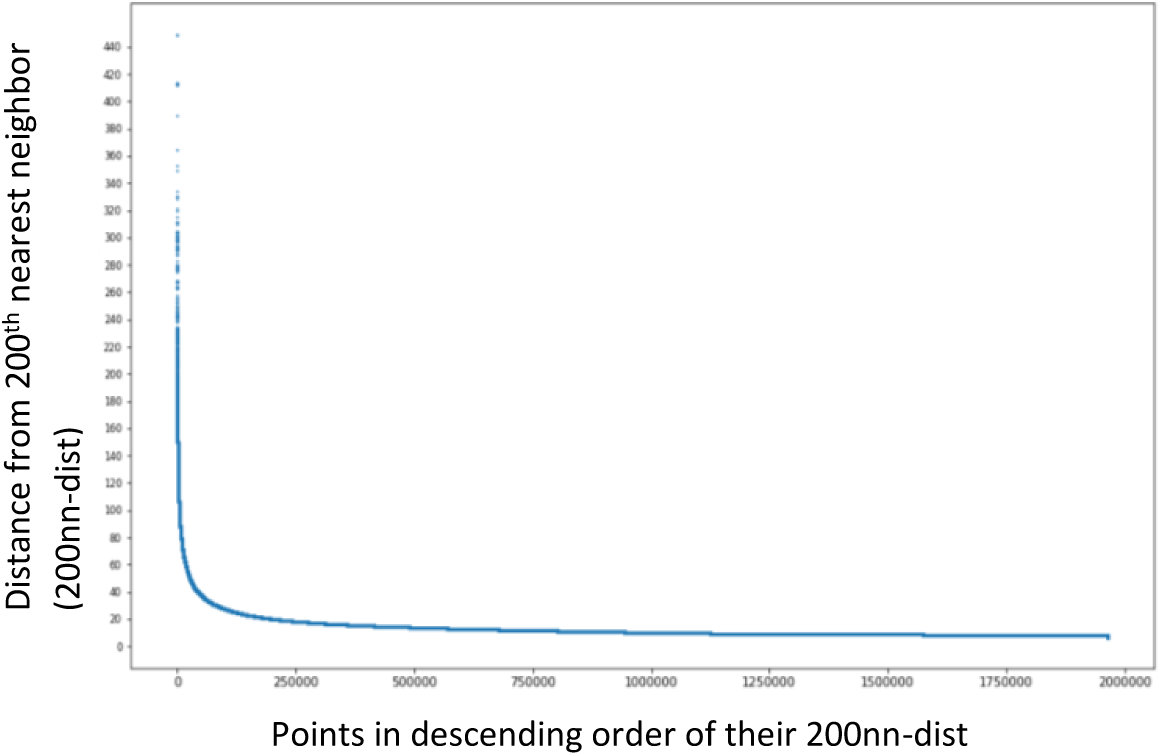
HMM+ pixels sorted by distance from 200^th^ nearest neighbor. This plot was used to visualize the distribution of 200^th^ nearest neighbor distances across HMM+ pixels.

**Supplementary Figure 8.**
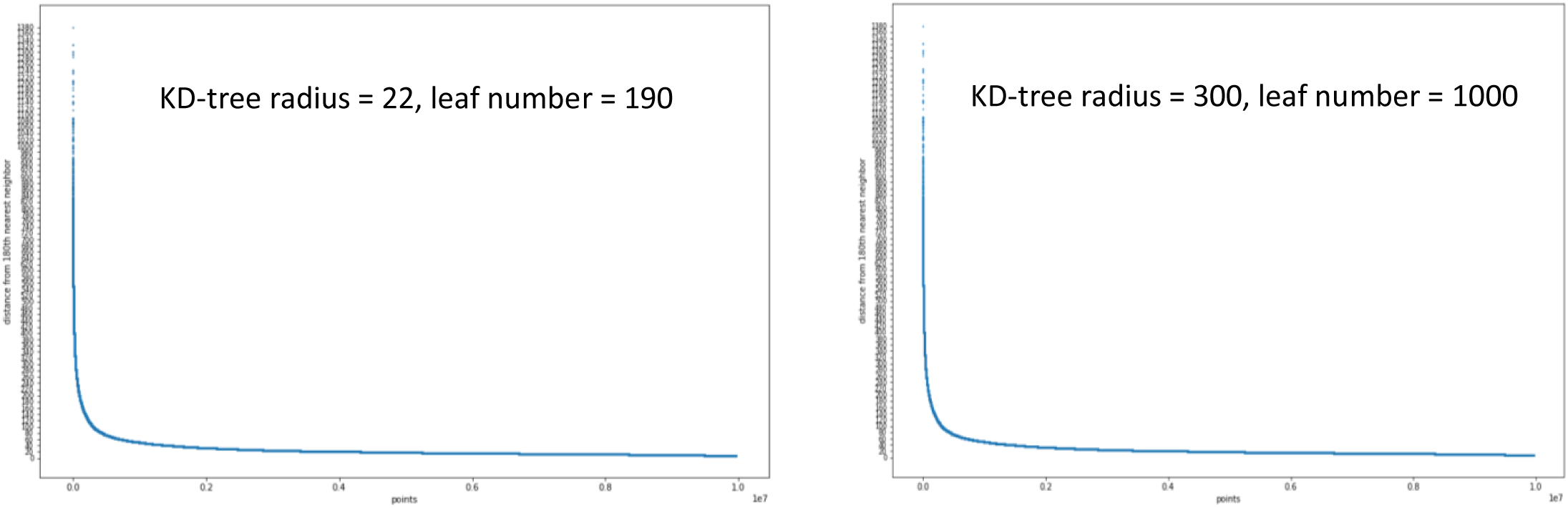
Example K-dist graphs.

We used the knee point of the first k-dist graph (at minPts = 200; Supplementary Figure 6) to estimate the optimal values for eps to be 20-30. We then ran DBSCAN on a grid of parameters and quantified the number of clusters labeled under each set of parameters. Three observers then annotated three sets of clips corresponding to minPts and eps values (indicated by the red outlines in Supplementary Figure 7). After comprehensive review, we decided the eps = 18 and minPts = 170 best reflected true sand change clusters.

**Supplementary Figure 9.**
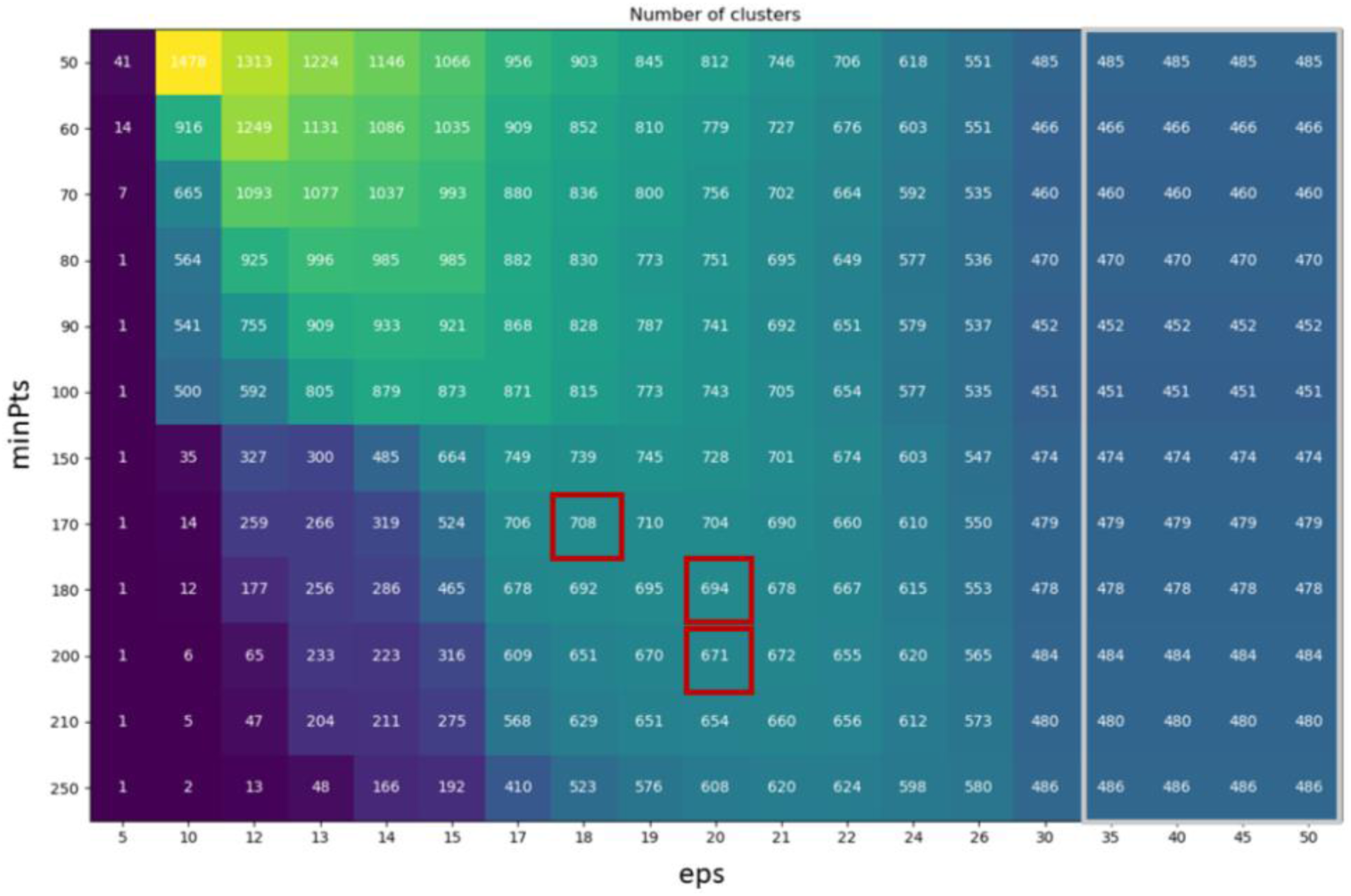
Number of identified clusters under different values for minPts and eps. This plot shows the number of identified clusters from segment of video data using different values for minPts and eps. Red boxes indicate values at which trained observers reviewed video clips of sand change clusters to identify optimal values for minPts and eps.

#### Nearest Neighbor KD-tree treeR/neighborR and leaf size

1. treeR and neighborR are equivalent parameters for constructing KD-trees (Pedregosa, Varoquaux et al. 2011). Within a radius around each point, all distances between this point and other points are calculated. DBSCAN queries the distances within eps (eps=18 in our analysis) for each point, so the treeR/neighborR ≥ eps. We set this parameter to 22 to prepare the distance matrix for DBSCAN with eps <= 22.
2. leaf_size: this parameter is a threshold below which the calculation switches from traversing tree to brute-force. For small data sets (N less than 30 or so), brute force algorithms can be more efficient than a tree-based approach. Changing leaf_size will not affect the results of a query, but can significantly impact the speed of a query and the memory required to store the constructed tree as seen in (Pedregosa, Varoquaux et al. 2011) and here: https://jakevdp.github.io/blog/2013/04/29/benchmarking-nearest-neighbor-searches-in-python/#Scaling-with-Leaf-Size. We set leaf_size above minPts 170 (leaf_size=190).

#### “Timescale”

Since DBSCAN uses one radius to search clusters in all dimensions, we scaled the time dimension so that the temporal lengths of events were similar in magnitude to their spatial width, such that events were, in general, roughly spherical in 3D. From watching the video we observed that the duration of sand change events was < 5 seconds, and their spatial widths were < 60 pixels; so the time dimension (on frame/second) was scaled by 10x.

### Behavioral definitions for manual annotation

#### Bower scoop

subject male collects sand into its mouth during bower construction.

#### Bower spit

subject male expels sand into its mouth during bower construction.

#### Bower spit

multiple bower scoops, multiple bower spits, and/or a bower scoop and bower spit are expressed by the subject male within the same video clip.

#### Feeding scoop

fish collects sand into its mouth during feeding.

#### Feeding spit

fish expels sand into its mouth during feeding.

#### Feeding multiple

multiple feeding scoops and/or spits are expressed by a fish within the same video clip.

#### Spawn/quiver

the subject male rapidly vibrates his body left to right while simultaneously circling, often but not necessarily with a female in frame. The male’s body is typically arched left to right, with his anal fin (egg spots) displayed directly in front of the female. When the female is present she is often circling as well.

#### Sand dropping

A fish expels or releases sand from the mouth either while high in the water (after which the sand sprinkles down through the water until it eventually lands), or release of sand upon initiation of a rapid burst of swimming (typically chasing or being chased). A more rare subset of sand dropping includes filtering sand through the operculum while swimming, typically during feeding.

#### Other

Changes to the sand caused by any other fish activity not described above, often as a result of swiping of the fin or rubbing of the ventral surface of the body along the sand during performance of other behaviors. More rare cases included instances in which two fish both perform behaviors in the same clip but the sand change was designated as a single cluster.

#### Shadow/reflection

Other changes that are not caused by fish manipulating or changing sand, most commonly reflections of activity in the aquarium glass and shadows cast by a stationary or very slow-moving fish, or in rare instances food, feces, or other debris settling on the sand surface.

**Supplementary Figure 10.**
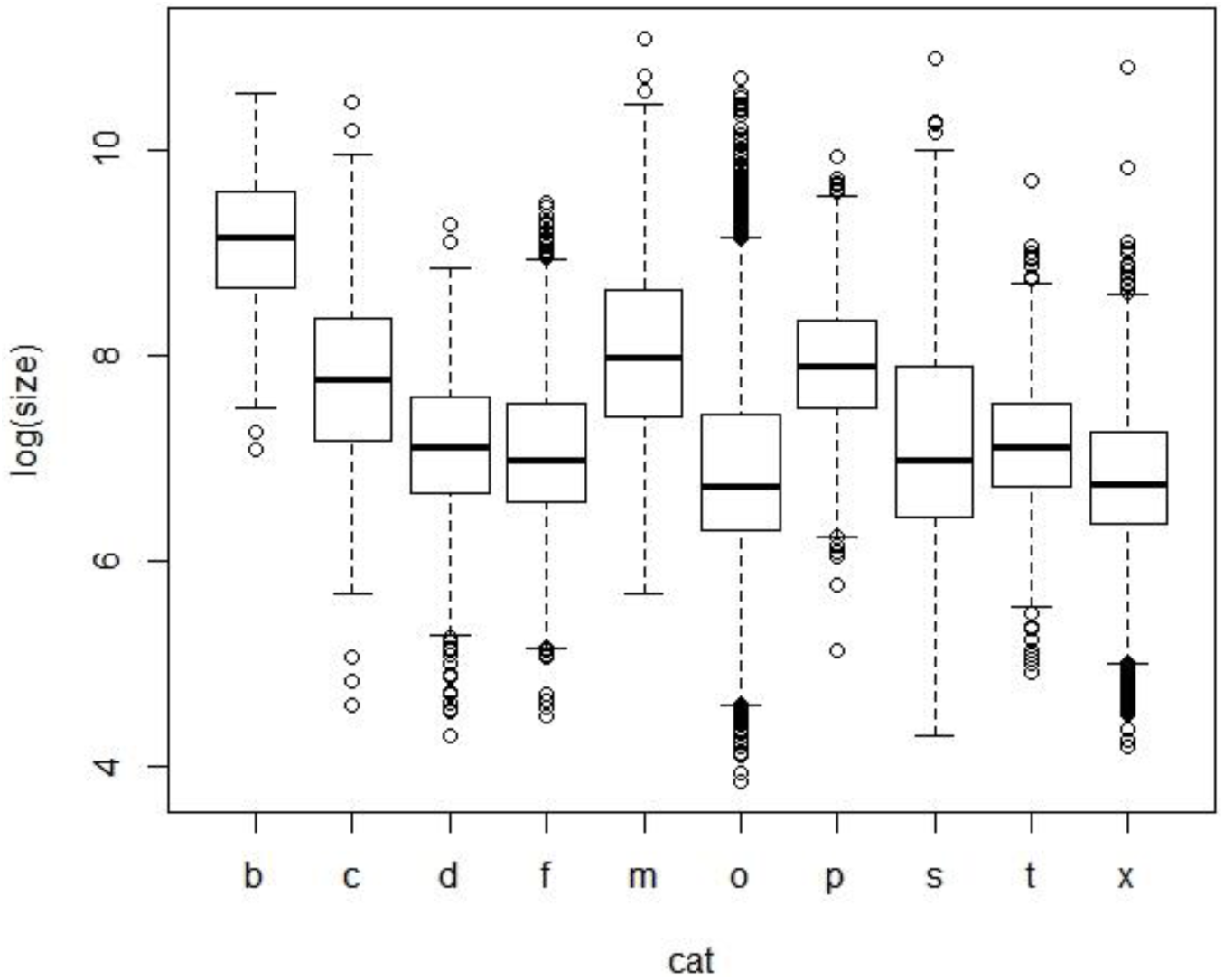
Cluster size by category. The relative pixel sizes (log) of clusters assigned to different categories (e.g. feeding, scoops and spits during bower construction, and quivering) were analyzed to determine their predictive value. We found significant differences in the size of clusters across eight behavioral and two other categories (Kruskal-Wallis rank sum test, χ=4223.4, p<2.0×10^−16^). The vast majority of pairwise comparisons were significant after correcting for multiple comparisons (Dunn’s test, adjusted p<0.05 for 40/45 pairwise comparisons between categories). However, the distributions of cluster size by category were highly overlapping and therefore cluster size alone was not sufficient for linking sand change events to different behaviors. Abbreviations: b = multiple bower events, c = bower scoop, d = sand dropping, f = feeding scoop, m = multiple feeding events, o = “other”, p = bower spit, s = quiver/spawn, t = feeding spit, x = shadow/reflection.

**Supplementary Figure 11.**
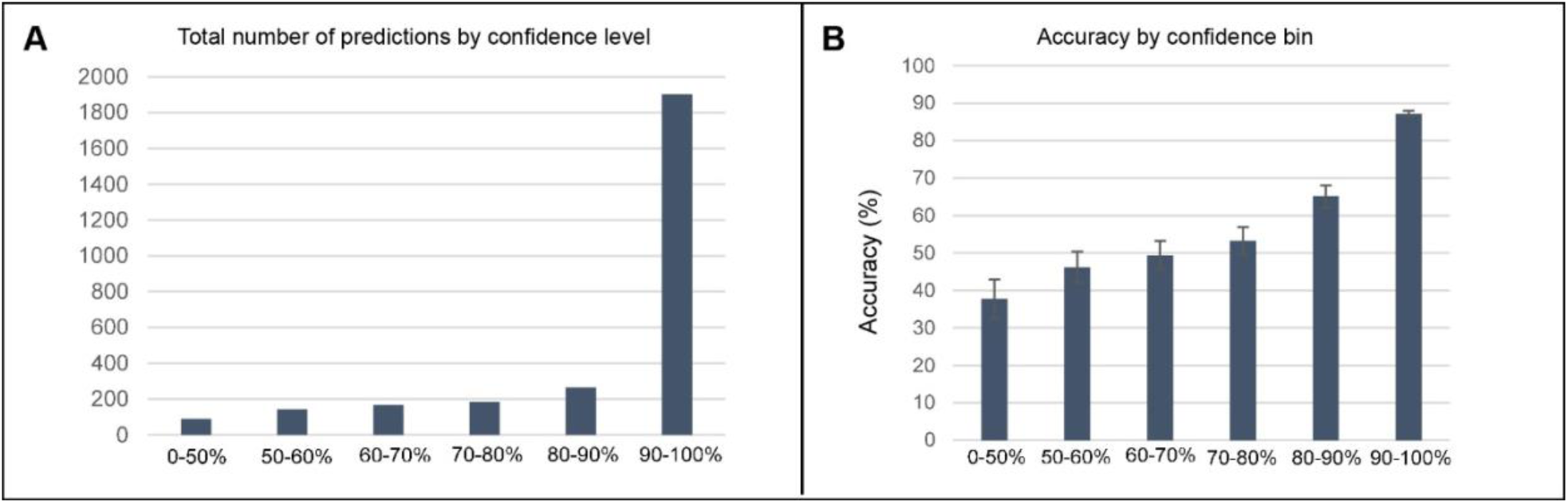
Relationships between confidence and accuracy for predictions by 3D ResNet for action recognition. The large majority of predictions (69%) were associated with high confidence scores (90-100%; A). High confidence (>90%) predictions tended to be more correct than low confidence predictions (B).

**Supplementary Figure 12.**
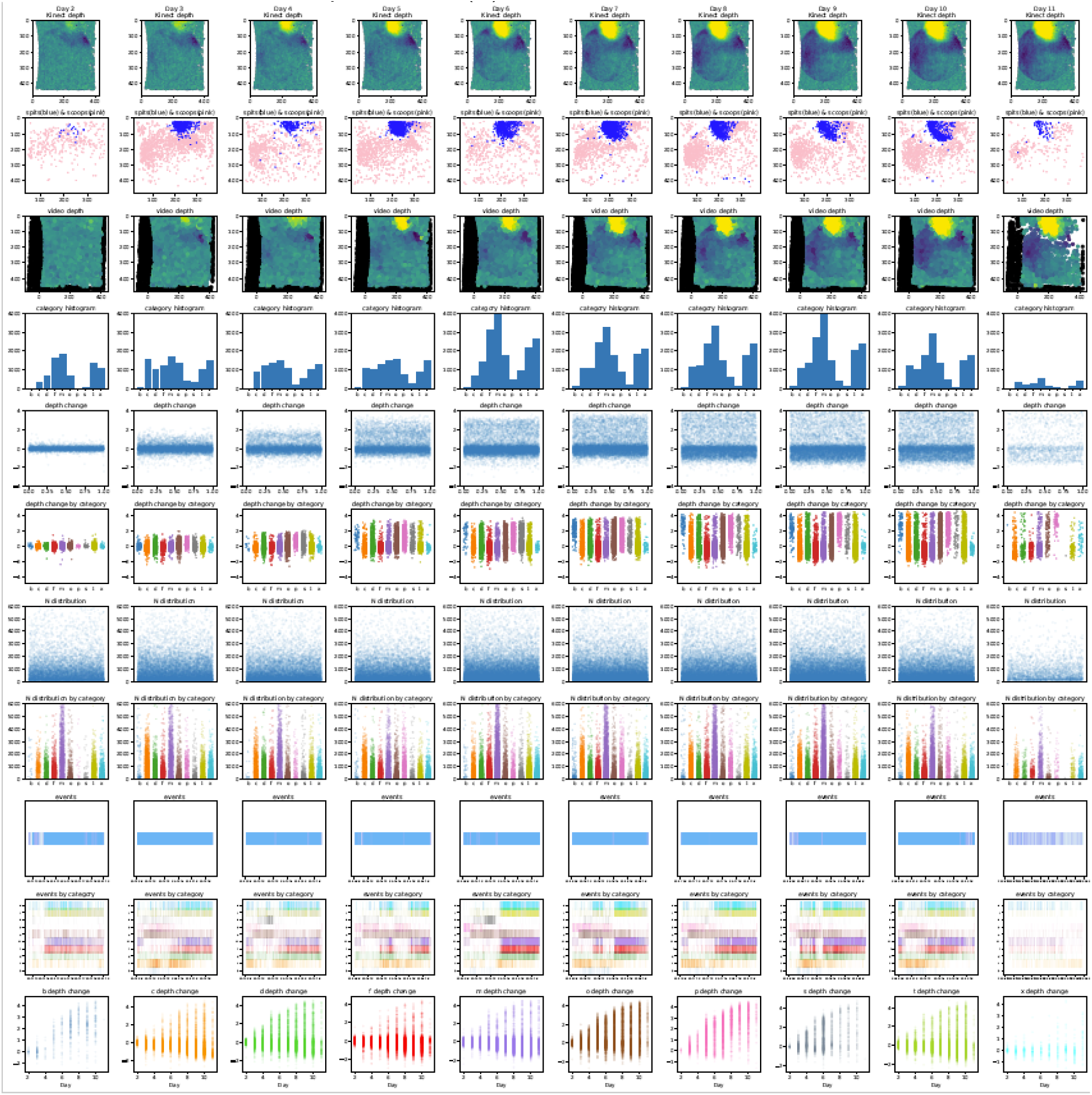
Example output following analysis of registered 3D ResNet-predicted behavioral events with depth sensing data across full trials. Visualization of behavioral analyses of a representative *Mchenga conophoros* trial. Depth change by day as measured by the Kinect across the full trial (first row). Spatial location of all 3D ResNet-predicted bower scoop (pink) and bower spit (blue) events across the full trial (second row). Depth of all behavioral events by day across the full trial (third row). Number of events across categories by day (fourth row). Depth change at locations of all behavioral events by day (fifth row). Depth change at locations of all behavioral events by category across days (sixth row). Pixel size of all sand change clusters by day (seventh row). Pixel size of all behavioral events by category across days (eighth row). Temporal distribution of all behavioral events by day (ninth row). Temporal distribution of all behavioral events by category across days (tenth row). Sand surface height at location of each behavioral event across days (consecutive data columns within each plot), by category (each consecutive plot represents a different behavioral category; eleventh row).

**Supplementary Table 1.** Actual and expected overlap by species and cross. Sample sizes for each species and cross used for spatial repeatability analysis, with mean (± S.E.) observed overlap and expected overlap between repeatability trials. These metrics are also shown for analysis of all subjects pooled together (bottom row).

| Species/<br>cross | N | Actual overlap %<br>(mean ± SE) | Expected overlap %<br>(mean ± SE) |
| --- | --- | --- | --- |
| CV | 5 | 25.9 ± 6.32 | 1.43 ± 0.76 |
| TI | 2 | 29.8 ± 17.40 | 2.03 ± 0.78 |
| MC | 4 | 14.3 ± 6.86 | 3.27 ± 1.65 |
| MCxCVF<sub>1</sub> | 1 | 51.4 | 6.39 |
| TIxMCF<sub>1</sub> | 2 | 14.7 ± 2.51 | 1.93 ± 1.05 |
| Pooled | 14 | 23.4 ± 4.30 | 2.5 ± 0.64 |

**Video Figure 1. CNN-predicted behavioral events by species, category, and test subject.** Subset of high confidence (>90%) predictions for each behavioral category (rows) by subject (columns), ordered from left to right by species and hybrid cross (CV, TI, MC, MCxCV F_1_ hybrid).

